# Coral anthozoan-specific opsins employ a novel chloride counterion for spectral tuning

**DOI:** 10.1101/2024.12.12.628111

**Authors:** Yusuke Sakai, Saumik Sen, Tomohiro Sugihara, Yukiya Kakeyama, Makoto Iwasaki, Gebhard F.X. Schertler, Xavier Deupi, Mitsumasa Koyanagi, Akihisa Terakita

**Author notes:** Corresponding authors: Xavier Deupi Mitsumasa Koyanagi Akihisa Terakita.

## Abstract

Animal opsins are G protein coupled receptors that have evolved to sense light by covalently binding a retinal chromophore via a protonated (positively charged) Schiff base. A negatively charged amino acid in the opsin, acting as a counterion, stabilizes the proton on the Schiff base, which is essential for sensitivity to visible light. In this study, we investigate the spectroscopic properties of a unique class of opsins from a reef-building coral belonging to the anthozoan-specific opsin II group (ASO-II opsins), which intriguingly lack a counterion residue at any of established sites. Our findings reveal that, unlike other known animal opsins, the protonated state of the Schiff base in visible light-sensitive ASO-II opsins is highly dependent on exogenously supplied chloride ions (Cl^−^). By using structural modelling and QM/MM calculations to interpret spectroscopy data, we conclude that, in the dark state, ASO-II opsins employ environmental Cl^−^ as their native counterion, while a nearby polar residue, Glu292 in its protonated neutral form, facilitates Cl^−^ binding. In contrast, Glu292 plays a crucial role in maintaining the protonation state of the Schiff base in the light-activated protein, serving as the counterion in the photoproduct. Furthermore, Glu292 is involved in G protein activation of the ASO-II opsin, suggesting that this novel counterion system coordinates multiple functional properties.

## Introduction

Animals sense light by using opsins, photosensitive proteins belonging to the large family of G protein-coupled receptors (GPCRs). These proteins have a seven-transmembrane helix structure and bind to a retinal chromophore to form a light-sensitive pigment. Opsins are present in the genomes of all eumetazoans (i.e., all animal lineages except sponges), and based on their phylogenetic relationships, they can be classified into eight groups with distinctive properties: vertebrate visual/non-visual opsins, opn3/TMT opsins, invertebrate Go-coupled opsins, cnidarian Gs-coupled opsins (cnidopsins), neuropsins (opn5), Gq-coupled visual pigments/melanopsins (opn4), peropsins, and retinochrome/RGR (Koyanagi and Terakita, 2014). Such diversity possibly underlies the diversification of light-dependent physiologies in animals. Furthermore, this diversity also provides a range of potential optogenetic tools to manipulate intracellular G protein-mediated signaling (Koyanagi and Terakita, 2014).

Reef-building corals and sea anemones belong to the subphylum Anthozoa, which together with the subphylum Medusozoa constitute the phylum Cnidaria. Cnidarian animals possess multiple opsins categorized as part of the cnidarian Gs-coupled opsin group (cnidopsins), which regulate light-dependent processes (Koyanagi et al., 2008). For example, a member of this group is expressed in the ciliary type visual cells of the box jellyfish lens eyes (Koyanagi et al., 2008; Kozmik et al., 2008).

Beyond these Gs-coupled cnidopsins, anthozoan animals have opsins that are phylogenetically distinct from the other known eight groups and are found exclusively in anthozoans (Feuda et al., 2012; Suga et al., 2008). These anthozoan-specific opsins (ASO) can be further classified into two groups, ASO-I and ASO-II (Gornik et al., 2020; Hering and Mayer, 2014; Mason et al., 2023; Picciani et al., 2018; Ramirez et al., 2016). Gornik et al. (2021) proposed that both ASO-I and ASO-II were present in the last common ancestor of Anthozoa and Medusozoa, but were lost secondarily in the Medusozoa lineage (Gornik et al., 2020). While it has been reported that both ASO-I and ASO-II are expressed in multiple tissues of sea anemones (Gornik et al., 2020; Suga et al., 2008) and corals (Levy et al., 2021), there is still a limited understanding of their molecular characteristics and physiological functions.

The members of the ASO-II group are not only phylogenetically unique but also display interesting features in their amino acid sequences. For instance, several of these opsins lack an amino acid residue conserved among typical opsins that is crucial for absorption of visible light (Gornik et al., 2020; Mason et al., 2023). While free retinal in solution has its absorption maximum (λ_max_) in the ultraviolet (UV), this shifts to visible light when retinal is bound to a lysine residue in the transmembrane bundle of the opsin (usually at Lys296, numbering according to the bovine rhodopsin sequence) through a protonated Schiff base to form the pigment. Such protonated Schiff base is necessary to achieve sensitivity to visible light in opsin-based pigments (Pitt et al., 1955). However, the proton on the positively charged Schiff base is energetically unstable in the hydrophobic transmembrane environment. To stabilize this proton, a negatively charged residue, glutamic or aspartic acid, is situated near the Schiff base to act as a counterion. This counterion is essential for opsin-based pigments to absorb visible light, and the residues serving as the counterion are highly conserved across opsins (Nathans, 1990; Terakita, 2005; Terakita et al., 2012). To date, three experimentally confirmed sites for the counterion have been identified in animal opsins: 94 in helix 2 (Gerrard et al., 2018), 113 in helix 3 (Nathans, 1990; Sakai et al., 2022; Sakmar et al., 1989; Zhukovsky and Oprian, 1989) and 181 in extracellular loop 2 (Nagata et al., 2019; Terakita et al., 2004, 2000). Remarkably, some opsins belonging to the ASO-II group lack glutamic or aspartic acid at any of these established counterion positions (Gornik et al., 2020; Mason et al., 2023). This absence raises the question of whether these opsins can absorb visible light, and if so, by what mechanism.

In this study, we investigate the spectroscopic properties of opsins in the ASO-II group isolated from the reef-building coral *Acropora tenuis*. Absorption spectra reveal that this group includes opsins sensitive to both UV and visible light. We then focus on a particular visible light-sensitive opsin within the ASO-II group (Antho2a) by spectroscopically analyzing the protonated and deprotonated states of the Schiff base in the wild type and in single point mutants. By interpretating the spectroscopy data in the light of hybrid quantum mechanics/molecular mechanics (QM/MM) simulations, we demonstrate that a chloride anion (Cl^−^) serves as a counterion to the retinylidene Schiff base in animal opsins, specifically in visible light-sensitive opsins of the ASO-II group.

## Results

### Identification of *Acropora tenuis* opsins

We identified 17 opsins from the *Acropora tenuis* genome and transcriptome datasets by homology search, which included eight opsins in Gs-coupled cnidopsin group, one opsin in the ASO-I group, and eight opsins in the ASO-II group (Fig. S1; Fig. 1A). Full-length cDNAs of seven out of the eight opsins in the ASO-II group were isolated and cloned from adult or larval tissues of the coral (highlighted by bold letters in Fig. 1A). We failed to amplify one opsin in the ASO-II group (gene model ID in the OIST Marine Genomics Unit Genome Project (Shinzato et al., 2020): aten_s0263.g14) by RT-PCR possibly because of its little mRNA expression level. Amino acid sequence alignment shows that all the seven *A. tenuis* opsins in the ASO-II group lack a glutamic or aspartic acid at the established counterion positions 94, 113 or 181 (Fig. 1B; Fig. S2). These opsins also have no E(D)RY motif at the cytoplasmic end of helix 3 (Fig. S2), which is conserved throughout most class A GPCRs (Hofmann et al., 2009).

**Fig. 1.**
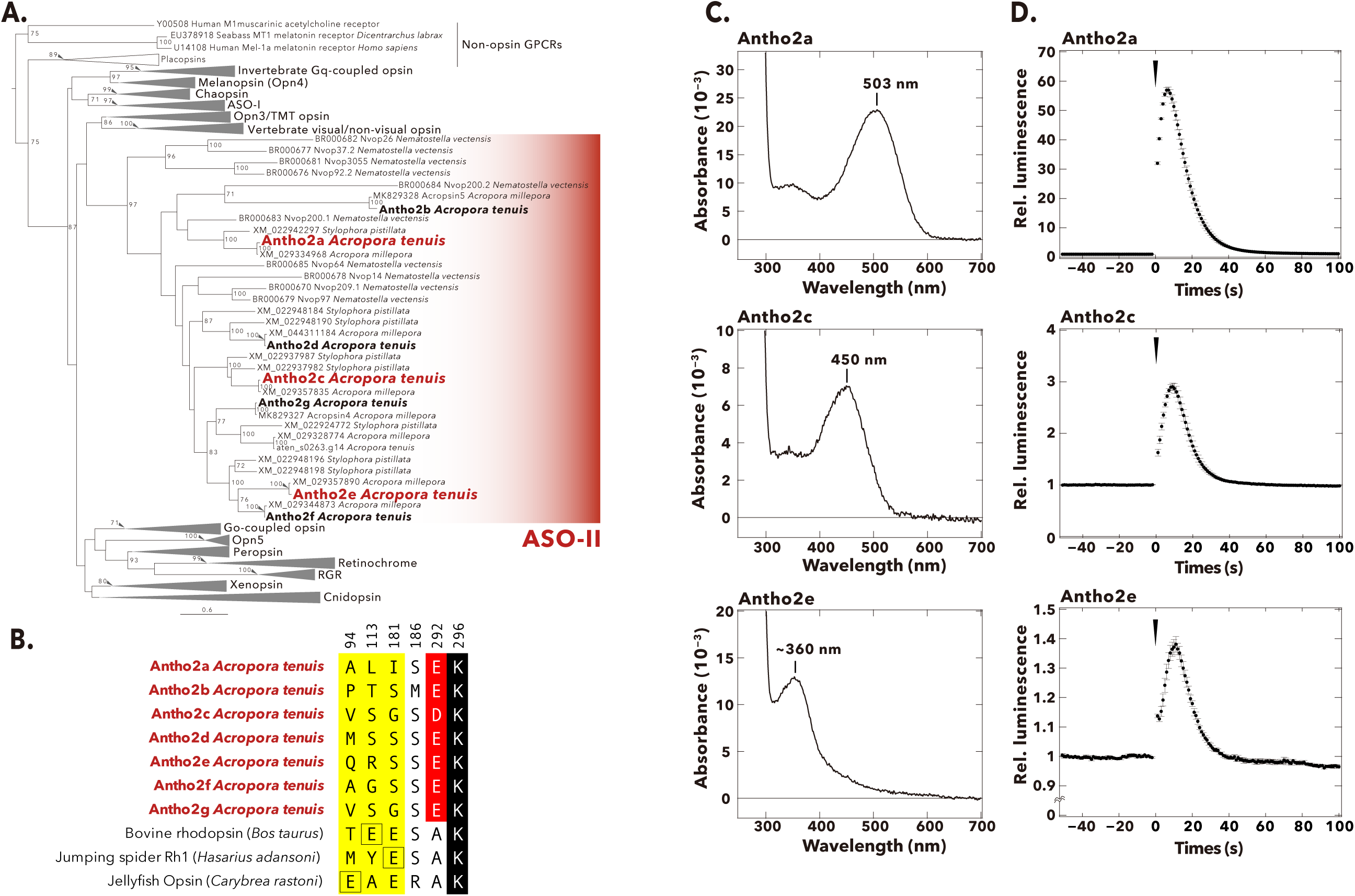
Phylogenetic tree, selected amino acid residues, absorption spectra, and light-induced Ca^2+^ responses of *Acropora tenuis* opsins belonging to the ASO-II group. **(A)** Maximum-likelihood (ML) tree of animal opsins including *A. tenuis* opsins in the ASO-II group. Seven opsins in the ASO-II group that were identified and cloned from *A. tenuis* in this study are shown in bold and the three members for which we obtained absorption spectra are highlighted in red. Numbers at the nodes represent support values of each ML branch estimated by 1000 bootstrap samplings (≥ 70% are indicated). Scale bar = 0.6 substitutions per site. All branches and support values are provided in Fig. S1. **(B)** Selected residues near the Schiff base in opsins of the ASO-II group and other animal opsins. Animal opsins typically have an acidic residue acting as counterion at one of three established sites (yellow): E94 (e.g., jellyfish opsin), E113 (e.g., bovine rhodopsin), or E181 (e.g., jumping spider Rh1). Remarkably, opsins in the ASO-II group lack an acidic residue at any of these positions, but instead feature an acidic residue at position 292 (red). The retinal-binding lysine, Lys296, is shown in black. A more detailed sequence alignment is provided in Fig. S2. Residues are numbered according to bovine rhodopsin. **(C)** Absorption spectra in the dark of three *A. tenuis* opsins in the ASO-II group (Antho2a, Antho2c and Antho2e). The absorption spectra were measured at 0°C in 140 mM NaCl at pH 6.5. The number in each graph shows the λ_max_ value. **(D)** Results of the aequorin-based bioluminescent reporter assay for monitoring light-induced changes in Ca^2+^ in HEK293S cells expressing the same three opsins in the ASO-II group as in panel C. In each graph, luminescence values were normalized to the baseline. Black circles with error bars indicate the means ± SEMs (n = 3) of the measured relative luminescence. Black arrowheads at time 0 indicate the timing of one-minute irradiation with green (495 nm; for Antho2a and Antho2c) or UV (395 nm; for Antho2e) light.

### Absorption spectra of *Acropora tenuis* opsins in the ASO-II group

We expressed seven members of the ASO-II group in COS-1 cells and purified their recombinant pigments in detergent-solubilized conditions. We successfully obtained the absorption spectra of three (Antho2a, Antho2c and Antho2e) out of the seven members, which showed that Antho2a and Antho2c are visible light-sensitive opsins having with λ_max_ at 503 nm and 450 nm, respectively, whereas Antho2e is a UV-sensitive opsin with λ_max_ at ∼360 nm (Fig. 1C). We have previously reported that one opsin in the ASO-II group, acropsin 4 of the coral *Acropora millepora*, induces a light-dependent elevation of intracellular Ca^2+^ levels (Mason et al., 2023). Here, we showed that Antho2a, Antho2c and Antho2e evoked a similar light-dependent increase of Ca^2+^ levels in HEK293S cells (Fig. 1D).

### Search for the counterion in *Acropora tenuis* opsins of the ASO-II group

Antho2a and Antho2c form visible light-sensitive pigments in the dark (Fig. 1C) despite the lack of a negatively charged counterion at any of the established positions (Fig. 1B; Fig. S2). To investigate how the protonated Schiff base is stabilized in these opsins, we studied in more detail Antho2a (λ_max_ = 503 nm), as it could be expressed well in cultured cells and was stable in detergent-solubilized conditions (Fig. 1C).

(a) *Contribution of Glu292 to the absorption spectra of the dark state and photoproduct of Antho2a*

First, we searched for potential counterions at positions different from known established amino acid sites (91, 113, and 181) in the Antho2a sequence. Using the crystal structure of bovine rhodopsin (PDB ID: 1U19) as a template, we identified glutamic or aspartic acids located within 5Å of the Schiff base in Antho2a and other members in the ASO-II group. Notably, all *A. tenuis* opsins in this group contain a conserved glutamic/aspartic acid at position 292 (Fig. 1B; Fig. S2), positioned just one helix turn away from the retinal-binding residue Lys296. To determine whether Glu292 could function as the counterion in Antho2a, we mutated Glu292 to alanine and measured the absorption spectra. The absorption spectrum of the E292A mutant in the dark was nearly identical to that of wild type (Fig. 2A, B, curve 1), exhibiting a clear absorbance in the visible light region with only a slightly red-shifted λ_max_ (505 nm) at 140 mM NaCl and pH 6.5. This shows that a negative charge other than Glu292 may serve as a counterion in the dark state of wild type Antho2a.

**Fig. 2.**
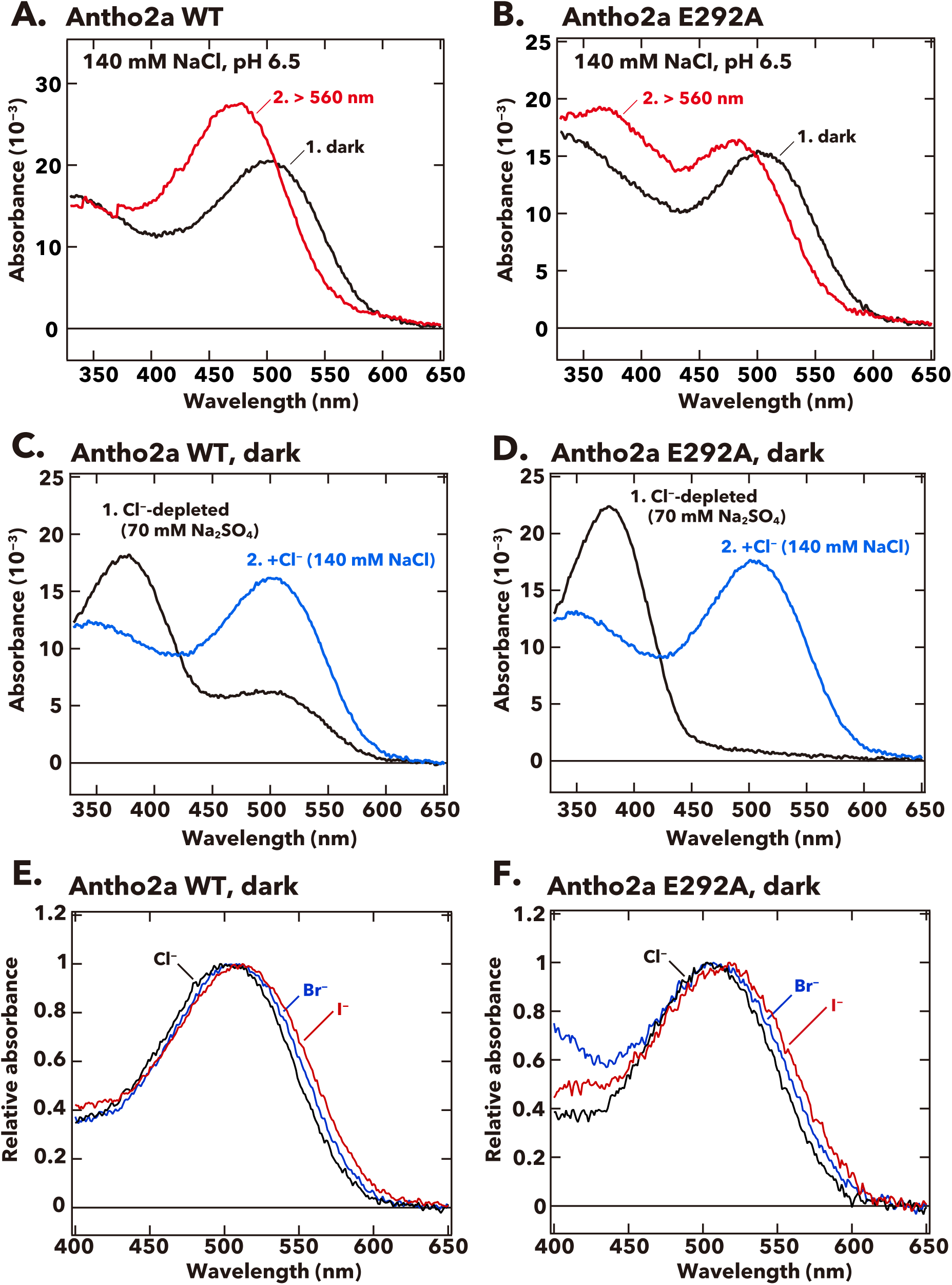
Absorption spectra of wild type and the E292A mutant of *A. tenuis* Antho2a. **(A, B)** Absorption spectra of the dark state (curve 1, black) and the photoproduct (curve 2, red) of the wild type (Antho2a WT, **A**) and the E292A mutant (Antho2a E292A, **B**) at 140 mM NaCl and pH 6.5. The samples were kept at 0°C during the spectroscopic measurements**. (C, D)** Absorption spectra of the dark state of Antho2a WT **(C)** and Antho2a E292A **(D)** prepared in Cl^−^-depleted conditions, before (curve 1, black) and after (curve 2, blue) adding Cl^−^ (see Materials and Methods for details). In the Cl^−^-depleted condition, the pigments were solubilized in 70 mM Na_2_SO_4_, which reportedly does not access to the Cl^−^ binding site in the chicken red-sensitive cone visual pigment iodosin (Shichida et al., 1990), to moderate protein denaturation. **(E, F)** Effect of halide anions on the absorption spectra of wild type Antho2a **(E)** and the Antho2a E292A mutant **(F)** at pH 6.5 and 0°C. The graphic shows the normalized absorption spectra of the pigments prepared in 140 mM NaCl (black curves), 140 mM NaBr (blue curves), and 140 mM NaI (red curves).

We next investigated the spectroscopic properties of the photoproduct of wild type Antho2a and the E292A mutant. Upon irradiation of wild type Antho2a with orange light, the λ_max_ shifted from 503 nm in the dark to 476 nm in the photoproduct (Fig. 2A, curve 2; Fig. S3A, curves 2 and 3). This shift is due to the photoisomerization of the 11-*cis* retinal chromophore to its all-*trans* form, converting almost 100% of the dark state to the photoproduct (Fig. S3B). The photoproduct remained stable for at least 5 minutes (Fig. S3A, curves 2 and 3) but did not revert to the original dark state upon subsequent irradiation (Fig. S3A and C). Instead, it underwent gradual decay accompanied by retinal release over time (Fig. S3D–G). These findings indicate that purified Antho2a is neither strictly bleach resistant nor bistable (see also Fig. S3 legend). We also observed that the protonated photoproduct decayed more rapidly at pH 8.0 (Fig. S3H) than at pH 6.5 (Fig. 3A, D, E). In contrast to the dark state, the photoproduct of the E292A mutant displayed two distinct absorption peaks in UV and visible light regions, at ∼370 nm and 476 nm, respectively (Fig. 2B, curve 2). This suggests that the E292A mutation causes UV-light absorption due to a deprotonated Schiff base in the photoproduct.

**Fig. 3.**
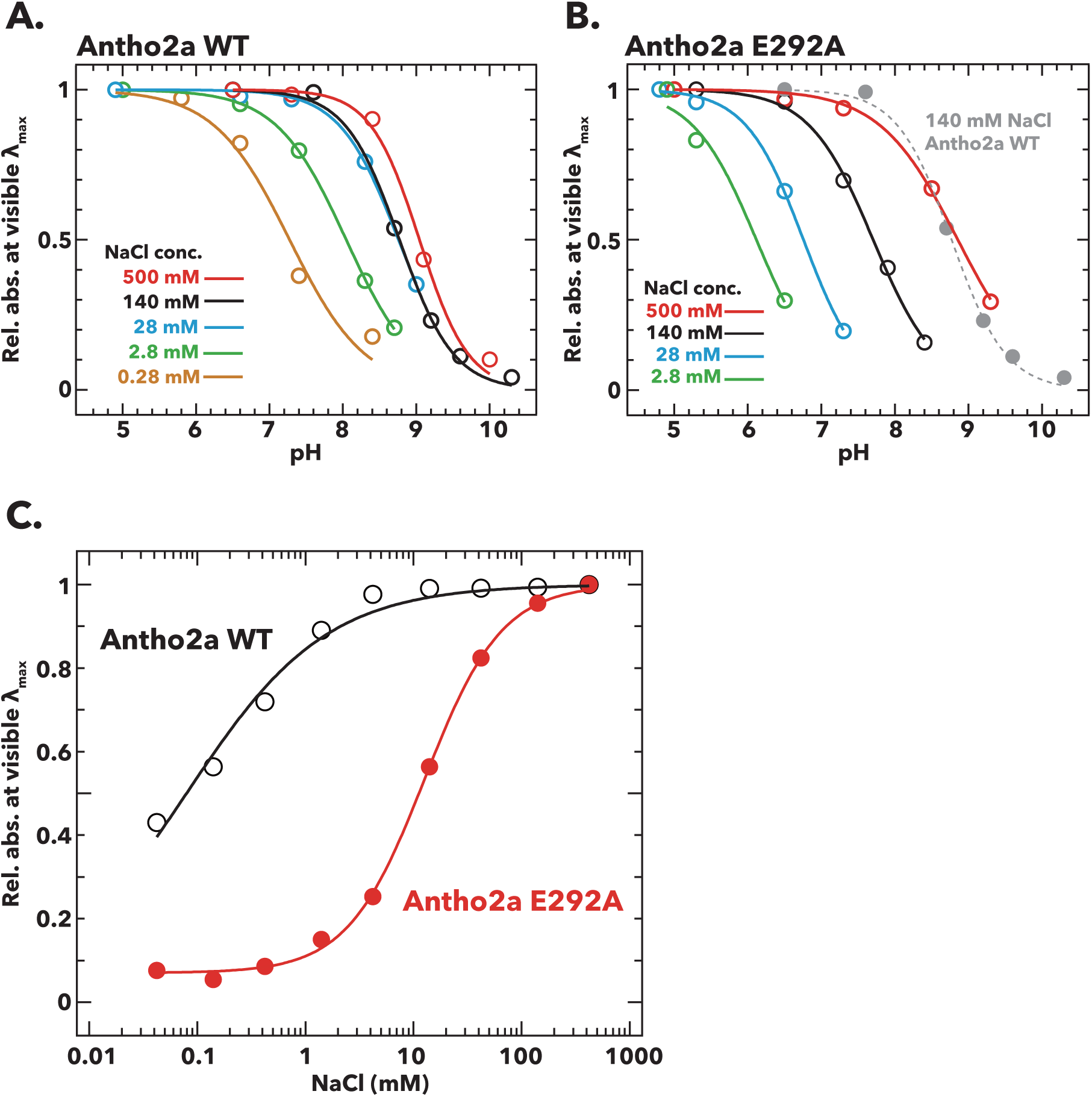
Effects of pH and Cl^−^ concentration on the absorption spectra of the dark states of wild type Antho2a and the Antho2a E292A mutant. **(A, B)** Changes in the absorbance at λ_max_ as a function of pH for **(A)** wild type Antho2a and **(B)** the E292A mutant at different Cl^−^ concentrations. The absorbance values at “visible λ_max_” (mean absorbance at 503 ± 5 nm for the wild type and 505 ± 5 nm for the E292A mutant, respectively) were normalized for each Cl^−^ concentration to those at the lowest pH, in which the Schiff base is assumed to be fully protonated (“Rel. abs. at visible λ_max_” in the y-axes). Solid and dashed lines represent sigmoid fits to the experimental data for each Cl^−^ concentration (indicated by different colors). The pH-dependent change of wild type Antho2a at 140mM NaCl is also shown in panel B (dotted grey line). The full absorption spectra used to generate these plots are provided in Fig. S6 (for wild type Antho2a) and Fig. S8 (for the E292A mutant). **(C)** Changes in the absorbance at λ_max_ for wild type Antho2a (black open circles) and the E292A mutant (red solid circles) as a function of Cl^−^ concentration. The absorbance values at visible λ_max_ were normalized to those at 500 mM NaCl for both the wild type and the E292A mutant. The lines in the graph were generated by fitting the Hill equation to the experimental data. The full absorption spectra used to generate these plots are provided in Fig. S9.

Additionally, altering the pH modified the ratio of absorbance between the ∼370 nm and 476 nm peaks in the E292A mutant (Fig. S4B, curves 2), with the UV-peak to the visible light-peak ratio increasing at higher pH levels (pH 7.4, Fig. S4B, curve 2). Conversely, the wild type did not exhibit an increase in UV absorbance under similar high pH condition (pH 7.5, Fig. S4A, curve 2). These results indicate that the Schiff base in the photoproduct of the Antho2a E292A mutant has a lower acid dissociation constant (p*K_a_*) than that of the wild type, suggesting that Glu292 acts as the counterion in the photoproduct of Antho2a.

We then further explored the nature of the counterion in the dark state of Antho2a. Previous studies have shown that in the bovine rhodopsin E113A and E113Q mutants, as well as in the retinochrome E181Q mutant (referred to as “counterion-less” mutants), halide ions like Cl^−^ can act as “surrogate” counterions to stabilize the proton on the Schiff base. Consequently, these counterion-less mutants can still absorb visible light in the presence of Cl^−^ (Nathans, 1990; Sakmar et al., 1991; Terakita et al., 2000). To assess the potential role of Cl^−^ as a surrogate counterion in the dark state of the Antho2a E292A mutant, we performed spectroscopic analyses under Cl^−^-depleted conditions (Fig. 2C, D). We observed that the λ_max_ of the E292A mutant shifted to the UV region (Fig. 2D, curve 1).

Unexpectedly, a similar shift in absorption to the UV region was also observed in the wild type under the Cl^−^-depleted condition (Fig. 2C, curve 1). These results indicate that, in the absence of Cl^−^, the Schiff base in both the wild type and the E292A dark states becomes deprotonated. The subsequent addition of Cl^−^ (final concentration: 140 mM NaCl) restored clear absorbance in the visible light region (Fig. 2C, D, curves 2), showing that Cl^−^ facilitates the protonation of the Schiff base of the dark state even in the wild type. In contrast, the photoproduct of the wild type exhibited no significant change in the ratio of UV to visible-light absorption peaks at pH 6.5 across NaCl concentrations from 0.28 mM to 800 mM (Fig. S5). The photoproduct of the wild type consistently absorbed visible light under these NaCl conditions (curves 2 in Fig. S5A, B), suggesting that Cl^−^ has little impact on the Schiff base p*K_a_* in the photoproduct of wild type Antho2a. However, the photoproduct of the E292A mutant exhibited a pH-dependent shift in the ratio of UV to visible-light absorption between pH 4.8 and pH 7.6, even at 800 mM NaCl, where the dark state predominantly absorbed visible light (Fig. S5C). This further supports that Glu292 serves as the counterion in the photoproduct of Antho2a.

(b) *Effect of halide anions on λ_max_ values of the dark state of Antho2a*

To obtain a further evidence supporting the Cl^−^ counterion in the dark state of Antho2a, we examined the impact of different halide anions on the absorption spectrum in the dark state of Antho2a, as observed in the bovine rhodopsin counterion-less mutant (Nathans, 1990; Sakmar et al., 1991). Antho2a readily absorbed visible light in the presence of bromide ion (Br^−^) and iodide ion (I^−^) as well as Cl^−^ and the λ_max_ of wild type Antho2a shifted depending on the halide solutions (503 nm in 140 mM NaCl; 506 nm in 140 mM NaBr; 511 nm in 140 mM NaI solutions; Fig. 2E). The E292A mutant showed a similar shift in λ_max_ (505 nm in 140 mM NaCl; 507 nm in 140 mM NaBr; 517 nm in 140 mM NaI solutions; Fig. 2F).

(c) *Effect of Cl^−^ concentration on the pK_a_ of the protonated Schiff base of Antho2a*

To further investigate the influence of Cl^−^ on the protonation state of the Schiff base in the dark state of Antho2a, we estimated the p*K_a_* of the Schiff base by measuring the pH-dependent changes in the absorption spectra of Antho2a at different Cl^−^ concentrations. The pH-dependent equilibrium between the visible (protonated Schiff base) and UV (deprotonated Schiff base) forms revealed that their ratio changes with Cl^−^ concentration (Fig. S6A–E). A plot of the changes in absorbance at λ_max_ against pH (Fig. 3A) shows that in wild type Antho2a, the p*K_a_* of the protonated Schiff base increases with higher Cl^−^ concentrations (7.3 at 0.28 mM NaCl, 8.0 at 2.8 mM, 8.8 at 28 mM, 8.8 at 140 mM, and 9.0 at 500 mM). We failed to determine the p*K_a_* at 0 mM NaCl, as the observed λ_max_ in acidic conditions (pH < 6.5) were shorter than expected in Antho2a (503 nm), suggesting that a normal pigment was not produced under these conditions (Fig. S7A). Similarly, the Cl^−^ concentration also affected the p*K_a_* of the protonated Schiff base in the E292A mutant (6.1 at 2.8 mM NaCl, 6.8 at 28 mM, 7.7 at 140 mM, and 8.9 at 500 mM) (Fig. 3B; Fig. S8). At 0 mM NaCl, the E292A mutant showed no visible light absorption, even under the most acidic conditions (pH 4.7), preventing the determination of its p*K_a_* (Fig. S7B). Notably, at low Cl^−^ concentrations (2.8 mM NaCl), the wild type exhibited a higher p*K_a_* than the E292A mutant (8.0 and 6.1, respectively). However, at 500 mM NaCl, the p*K_a_* of the E292A mutant and wild type were comparable (9.0 and 8.9, respectively; Fig. 3A, B). These results suggest that Cl^−^, rather than Glu292, serves as the counterion in the dark state of Antho2a, while Glu292 facilitates the protonation of the Schiff base by the Cl^−^ counterion.

(d) *Binding affinity of Cl^−^ to wild type Antho2a and the E292A mutant*

To evaluate the Cl^−^ binding affinities of both wild type Antho2a and the E292A mutant, we measured changes in their absorption spectra by gradually increasing Cl^−^ concentrations at pH 6.5 and estimated the Cl^−^ dissociation coefficients (*K_d_*). The relative absorbance in the visible region increased with higher Cl^−^ concentrations both in the wild type and the E292A mutant (Fig. S9A, B). By fitting the Hill equation to the experimental data (Fig. 3C), the dissociation constants (*K_d_*) of Cl^−^ were determined to be 0.079 ± 0.010 mM for the wild type Antho2a and 12.7 ± 0.519 mM for the E292A mutant. This significant increase in the *K_d_* value for the E292A mutant suggests that the Cl^−^ binding affinity is considerably reduced due to the mutation. Consequently, we suggest that while Glu292 does not act as a direct counterion, it plays a crucial role in facilitating Cl^−^ binding to Antho2a.

### Structural modelling and QM/MM calculations of the dark state of Antho2a

To gain a deeper understanding of the environment surrounding the retinylidene Schiff base in the dark state of Antho2a, we performed QM/MM-based structural modelling of both the wild type Antho2a (with Glu292 either neutral or negatively charged) and the E292A mutant. The QM/MM geometry optimization positioned the Cl^−^ ion close to the Schiff base (∼3Å) and near Glu292 (∼4.7Å), with Glu292 itself located in proximity to the Schiff base (∼3.3Å) (Fig. 4B). The chloride ion is also coordinated by two water molecules and the backbone of Cys187 which is part of a conserved disulfide bridge (Fig. S2). The retinylidene Schiff base region also includes polar (Ser186, Tyr91) and non-polar (Ala94, Leu113) residues (Fig. 4). To validate these models, we calculated the QM/MM vertical excitation energies of the ground state geometries (Table S1).

**Fig. 4.**
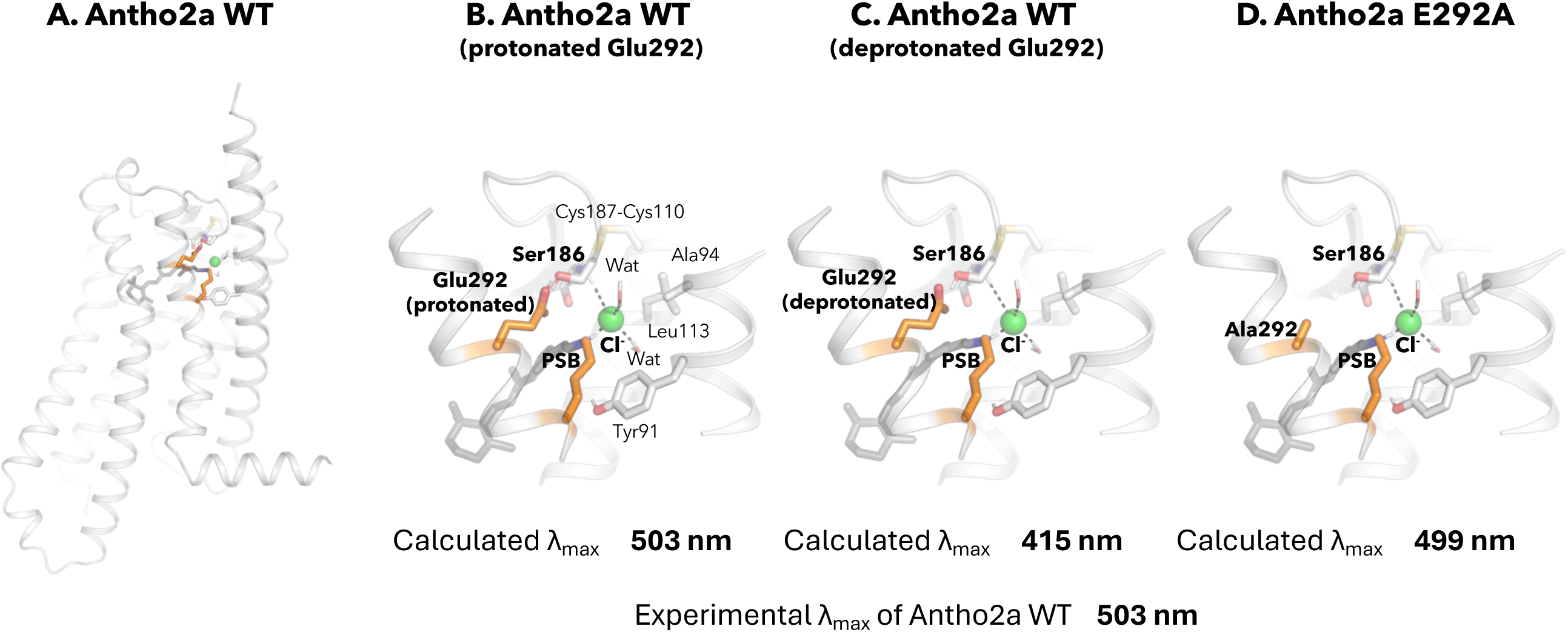
QM/MM structural model of wild type Antho2a in the dark state (A) and detailed views of the retinal binding pocket with a protonated (neutral) Glu292 (B), a deprotonated (negatively charged) Glu292 (C), and the E292A mutant (D). The retinal protonated Schiff base (PSB) and the binding pocket residues are shown as sticks (including polar hydrogens) and the Cl^−^ ion as a sphere with its coordination shown as dashes. “Wat” indicates a water molecule. Residues in the QM region are marked in bold.

For wild type Antho2a with a protonated neutral Glu292, the calculated λ_max_ using the CAM-B3LYP/cc-pVTZ level of theory was 503 nm (Fig. 4B), in good agreement with the experimentally observed value (503 nm; Fig. 2A). In contrast, the λ_max_ calculated with a deprotonated negatively charged Glu292 was blue-shifted to 415 nm (Fig. 4C), deviating significantly from the experimental value. Finally, the calculated λ_max_ for the E292A mutant was 499 nm (Fig. 4D), also in agreement with the experimental value (505 nm). To further substantiate these findings, we recalculated the excitation energies using the RI-ADC(2)/cc-pVTZ method. Although these λ_max_ values are blue-shifted compared to those calculated with the CAM-B3LYP method, they followed a similar trend. Both these computational methods have previously been employed to accurately calculate the excitation energies of rhodopsins (Church et al., 2021). These results strongly suggest that in the dark state of Antho2a, Glu292 is protonated and neutral at pH 6.5, and therefore, it does not function as the counterion.

### Effect of the Glu292 mutation on the function of the photoproduct

The spectroscopy data indicate that Glu292 is involved in stabilizing the protonated Schiff base by facilitating Cl**^−^** binding in the dark state and also serves as a counterion in the photoproduct. This suggests that Glu292 significantly contributes to the visible light absorption of Antho2a. To explore additional roles of Glu292 in Antho2a, we measured the light-induced Ca^2+^ response in cultured cells expressing wild type Antho2a or the E292A mutant. Notably, cells expressing wild type Antho2a showed a ∼30-fold increase in Ca^2+^ levels upon light irradiation (Fig. 5, solid black circles), whereas cells expressing the Antho2a E292A mutant showed a smaller Ca^2+^ elevation (< 5-fold increase) (Fig. 5, red open circles). This indicates that the peak Ca^2+^ response in cells expressing wild type Antho2a was approximately nine times greater than in cells expressing the E292A mutant. This result, along with the crucial role of Glu292 in Cl**^−^** binding in the dark state and as a counterion in the photoproduct, suggests that Glu292 plays also a role in G protein activation.

**Fig. 5.**
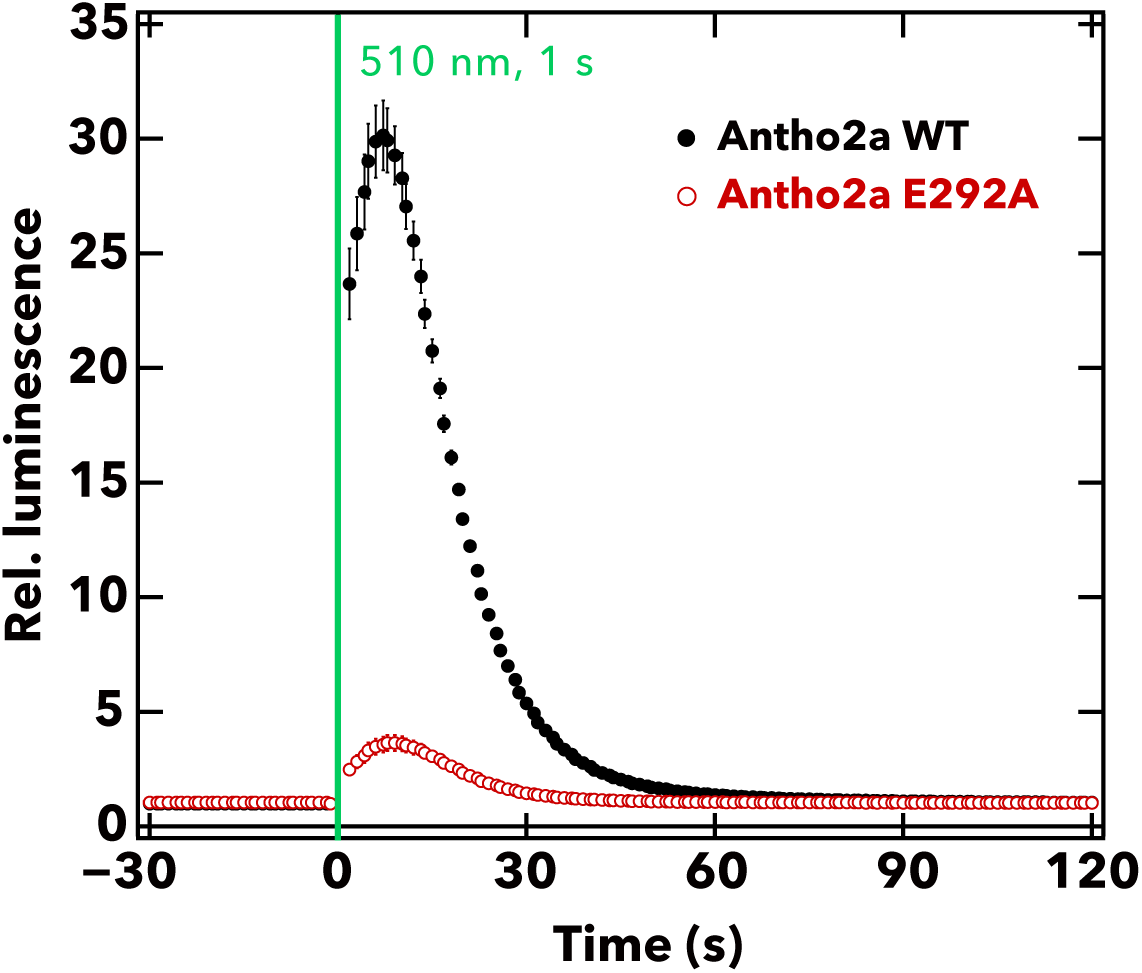
Comparison of the light-evoked intracellular Ca^2+^ levels between wild type Antho2a and the E292A mutant. The graph shows the mean ± S.E.M (n = 4) of the measured relative luminescence values (luminescence values normalized to the baseline) for wild type Antho2a (black) and the E292A mutant (red) at pH7.0. The green vertical line indicates the time of cell illumination with green light (510 nm, for 1 s, 1.65 × 10^15^ photons/cm^2^/s).

### Cl^−^-dependent changes in the absorption spectra of the dark states of Antho2c and Antho2e

We tested whether Cl^−^ concentration affects the p*K_a_* of the Schiff base in another visible light-sensitive opsin, Antho2c (λ_max_ = 450 nm, Fig. 1C). The pH-dependent equilibrium between UV- and visible-light absorbing forms was clearly observed at 0 mM or 0.093 mM NaCl, but not at 9.3 mM NaCl, where Antho2c stably absorbed visible light across the measured pH range (pH 4.8 to 7.2, Fig. 6A–C). Also, the ratio of UV to visible-light absorption increased with higher Cl^−^ concentrations at pH 6.5 (Fig. 6D). These results demonstrate that Cl^−^ serves as a counterion in the dark state of Antho2c, as it does in Antho2a. In contrast, wild type Antho2e continues to absorb UV light even at 1 M NaCl (Fig. 6E). Notably, Antho2e has an arginine at position 113, which corresponds to the counterion position in vertebrate visual opsins (Fig. S2). When this arginine is mutated to alanine (R113A), the mutant becomes sensitive to visible light (λ_max_ = ∼420 nm) in the presence of Cl^−^ (Fig. 6F), suggesting that Cl^−^ can serve as the counterion in the R113A mutant.

**Fig. 6.**
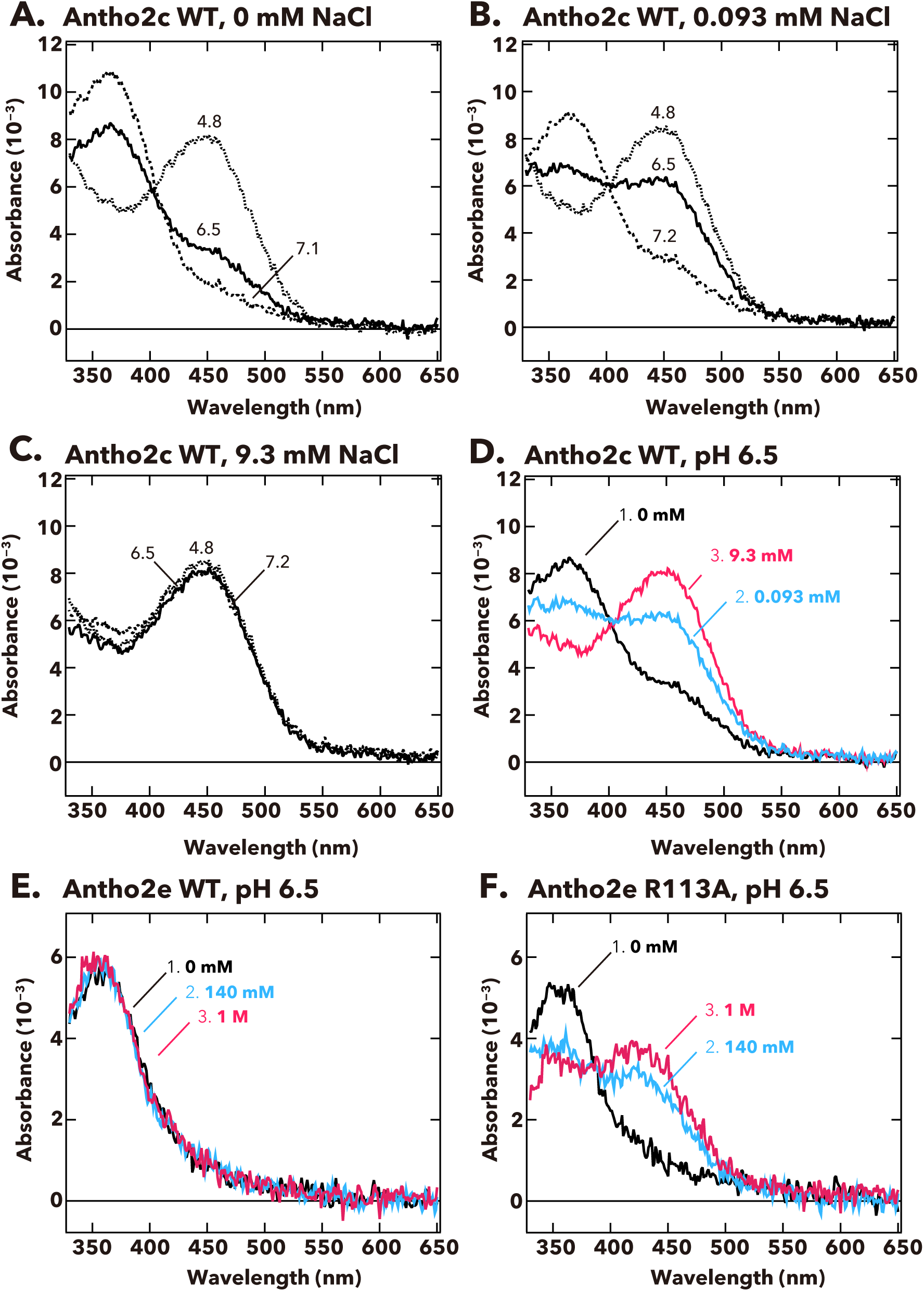
pH-dependent changes in the absorption spectra of Antho2c and Antho2e at different Cl^−^ concentrations at 0°C. **(A–C)** Absorption spectra of purified wild type Antho2c pigment at **(A)** 0 mM, **(B)** 0.093 mM, and **(C)** 9.3 mM NaCl concentrations. The corresponding pH values are indicated on each curve in the graphs. **(D)** Summary of the spectral changes for wild type Antho2c across different Cl^−^ concentrations at neutral pH (pH 6.5). **(E, F)** Absorption spectra of **(E)** wild type Antho2e (Antho2e WT) and **(F)** its R113A mutant (Antho2e R113A) at different Cl^−^ concentrations at pH 6.5 at 0°C. Each color indicates a different Cl^−^ concentration.

## Discussion

In this study, we reveal for the first time the spectral properties of opsins in the ASO-II group from the coral *Acropora tenuis*, showing that their sensitivity spans from UV to visible light. Opsins in this group have a highly conserved Glu292 residue near the Schiff base, which can potentially stabilize the proton on the Schiff base. Indeed, our results show that the p*K_a_* of the protonated Schiff base in the photoproduct (λ_max_ = 476 nm) of Antho2a is altered when Glu292 is substituted with alanine (Fig. 2A, B; Fig. S4), suggesting that Glu292 serves as the counterion of the photoproduct. Conversely, the dark state of Antho2a (λ_max_ = 503 nm) exhibits robust visible light absorption only in the presence of Cl^−^ at physiological pH, and the p*K_a_* of the protonated Schiff base changes with Cl^−^ concentration in both wild type Antho2a and the E292A mutant. Furthermore, the p*K_a_* for wild type and the E292A mutant are comparable in the presence of sufficient Cl^−^ (500 mM NaCl, Fig. 3A, B), supporting the conclusion that Cl^−^, and not Glu292, acts as the counterion of the dark state of *A. tenuis* Antho2a. We found that the type of halide anions in the solution has a small but noticeable effect on the λ_max_ values of the dark state of Antho2a. This is consistent with the effect observed in a counterion-less mutant of bovine rhodopsin, in which halide ions serve as surrogate counterions (Nathans, 1990; Sakmar et al., 1991). Similarly, our results align with earlier observations that the λ_max_ of a retinylidene Schiff base in solution increases with the ionic radius of halides acting as hydrogen bond acceptors (i.e., I^−^ > Br^−^ > Cl^−^) (Blatz et al., 1972). In contrast, the λ_max_ of halorhodopsin from *Natronobacterium pharaonic* does not clearly correlate with halide ionic radius (Scharf and Engelhard, 1994), as the halide ion in this case is not a hydrogen-bonding acceptor of the protonated Schiff base (Kouyama et al., 2010; Mizuno et al., 2018). Altogether, these findings support our hypothesis that in Antho2a, a solute halide ion forms a hydrogen bond with the Schiff base, thereby serving as the counterion in the dark state.

Moreover, QM/MM calculations for the dark state of Antho2a suggest that Glu292 is protonated and neutral, further supporting the hypothesis that Glu292 does not serve as the counterion in the dark state. However, unlike dark state, Cl^−^ has little to no effect on the visible light absorption of the photoproduct (Fig. S5). Therefore, we conclude that Cl^−^ and Glu292, respectively, act as counterions for the protonated Schiff base of the dark state and photoproduct of Antho2a. This represents a unique example of counterion switching from exogeneous anion to a specific amino acid residue upon light irradiation (Fig. S10).

Our spectroscopic data also showed that the other visual light sensitive opsin, Antho2c, exhibited the Cl^−^ dependency on the Schiff base p*K_a_* of the dark state, which suggesting that opsins in the ASO-II group may share a spectral tuning mechanism based on the Cl^−^ counterion. Interestingly, Antho2e had an arginine at position 113 and when it was mutated to alanine (R113A), the mutant showed sensitivity to visible light in the presence of Cl^−^ (Fig. 6F). We hypothesize that the positive charge of Arg113 disturbs interaction between the Cl^−^ and the protonated Schiff base or that it completely inhibits Cl^−^ binding, rendering Antho2e UV-sensitive.

It is noteworthy that although Cl^−^ has been reported to serve as a surrogate counterion in “counterion-less” mutants of animal opsins (such as E113Q in bovine rhodopsin and E181Q in retinochrome) (Nathans, 1990; Sakmar et al., 1991; Terakita et al., 2000), Antho2a is, to our knowledge, the first example in a wild type animal opsin that employs Cl^−^ as a counterion. Interestingly, within the microbial rhodopsin family, heliorhodopsin (TaHeR) incorporates a Cl⁻ into the Schiff base region under high Cl⁻ concentrations at pH 4.5. This Cl⁻ stabilizes the protonated retinal Schiff base when its primary counterion, E108, is neutralized (Besaw et al., 2022). The observed 18 nm red shift at low pH is consistent with E108 protonation. The TaHeR E108A mutant shows the same λ_max_ under high Cl⁻ concentrations, further supporting the role of Cl⁻ as a counterion. At pH 8, the Schiff base proton is stabilized by the negatively charged E108 (Shihoya et al., 2019).

The E292A mutation in Antho2a drastically decreases the Cl^−^ binding affinity in the dark state at pH 6.5 (*K_d_*: Antho2a WT = 0.079 mM, Antho2a E292A = 12.7 mM). Based on these results, supported by our QM/MM calculations of the dark state, we hypothesize that protonated Glu292 not only serves as a counterion of the photoproduct but also constitutes part of the Cl^−^ binding site in the dark state of Antho2a. In Cl^−^-pumping microbial rhodopsins, where Cl^−^ serves as a counterion, the Cl^−^ binding site near the Schiff base is typically formed by hydrogen-bonding network between Cl^−^, the protonated Schiff base, and key amino acids such as serine, threonine and glutamic/aspartic acid, and water molecules (Besaw et al., 2020; Hosaka et al., 2016; Kolbe et al., 2000). Additionally, the bovine rhodopsin double mutant E113Q/A292E exhibits a higher sensitivity to hydroxylamine than the wild type, indicating instability of the Schiff base and increased solvent accessibility (Tsutsui and Shichida, 2010). These observations suggest that Glu292 in Antho2a may also facilitate the accessibility of Cl^−^ ions to the Schiff base environment. To determine the precise nature of the Cl^−^ binding site and clarify the roles of Glu292 and other residues in the Cl^−^-dependent counterion system of Antho2a, detailed spectroscopic and structural experiments will be necessary.

We also found that cells expressing the Antho2a E292A mutant show a lower Ca^2+^ elevation upon light stimulation compared to the wild type (Fig. 5). The relative expression level of the E292A mutant of Antho2a was approximately 0.81 of the wild type (set as 1), as determined by comparing absorbances at λ_max_ for both pigments expressed and purified under identical conditions (Fig. S11A).

Additionally, the fraction of protonated pigment relative to the wild type (set as 1 at pH 6.5) was estimated to be 0.94 for the E292A mutant at pH 6.5, and 0.99 and 0.84 for the wild type and the E292A mutant at pH 7.0, respectively (Fig. 3A and B). Since pH 7.0 corresponds to the conditions used in the live cell Ca^2+^ assays, the effective amount of protonated pigment for the E292A mutant was approximately 73% of the wild type. Nevertheless, even after normalization for these differences, the Ca^2+^ response amplitude of the E292A mutant remained significantly lower (∼ 17% of wild type, compared to the observed 12% prior to normalization; Fig. 5 and Fig. S11B). These observations suggest that Glu292 serves not only as a counterion in the photoproduct but also plays an allosteric role in influencing G protein activation. It has been reported that the introduction of Glu or Asp at position 292 can affect various molecular properties of opsins. For instance, the A292E mutation in human rhodopsin has been shown to result in constitutively active apoproteins, leading to congenital night blindness (Dryja et al., 1993; Jin et al., 2003). These data suggest that Glu292 has the potential to influence functional properties, such as G protein activation. Several studies have reported that the counterion can affect diverse properties of opsins beyond their absorption spectra. For instance, in bovine rhodopsin, Glu113 is involved not only in visible light absorption in the dark state but also in the efficient activation of G proteins by the photoproduct (Terakita et al., 2004). It has been suggested that these pleiotropic functions of the counterion have been achieved through evolutionary modifications of the protein structure once the counterion was acquired (Terakita et al., 2004). This concept may similarly apply to Antho2a.

One of the notable features of the opsins in the ASO-II group that use Cl^−^ as a counterion is the relatively low p*K_a_* of the Schiff base (e.g., ∼9.0 for Antho2a in the dark state, Fig. 3A) compared to other animal opsins with a “regular” amino acid counterion (e.g., >16 for bovine rhodopsin, with Glu113 as a counterion (Steinberg et al., 1993) or 11 for jumping spider Rh1, with Glu181 as a counterion (Nagata et al., 2019)). We hypothesize that opsins in the ASO-II group, with lower Schiff base p*K_a_*, may change their spectral sensitivity and G protein activation profile within the physiological pH range of corals. For instance, these opsins could exhibit decreased sensitivity to visible and increased sensitivity to UV light under alkaline pH conditions. The extra- and intracellular pH environments in symbiotic cnidarians such as corals are spatially and temporally variable due to the photosynthesis of symbiotic algae (Barott et al., 2017). For example, pH values in the cytosol of algae-hosting cells (a group of endodermal cells with algal occupancy) in corals reportedly increase by approximately 0.5 pH units upon light treatment (Venn et al., 2009). The pH values in the gastrovascular cavity (coelenteron), which is in contact with endodermal cells, range from 6.6 to 8.5 (Agostini et al., 2012; Al-Horani et al., 2003; Cai et al., 2016) and increase in the presence of light, presumably due to the consumption of CO_2_ by photosynthesis (Barott et al., 2017). Additionally, alkalinization in extracellular region of calcifying cells reportedly increases ∼1.0 pH units from dark to light conditions reaching levels above pH 9.0 (Al-Horani et al., 2003). These pH changes, generally an increase due to photosynthetic activity, may result in variable light sensitivity of the “low p*K_a_*” opsins in the ASO-II group. Recent studies have reported that members of the ASO-II group may be associated with the symbiotic relationship between host anthozoans and symbiotic algae. For example, using the symbiotic sea anemone *Exaiptasia diaphana*, Gornik et al. (2021) showed that the mRNA levels of several opsins in the ASO-II group were higher in symbiotic adults than in apo-symbiotic ones (Gornik et al., 2020). In sea anemone, it has also been reported that behavioral responses to light differ between symbiotic and apo-symbiotic individuals (Foo et al., 2020; Kishimoto et al., 2023), with these responses potentially being driven by the activation of ASO-II opsins that are up-regulated in the presence of symbiotic algae. Moreover, single-cell RNA-seq analysis in the reef-building coral *Stylophora pistillata* has revealed that an opsin belonging to the ASO-II group is specifically expressed in algae-hosting cells (a group of endodermal cells with 50% algal occupancy) (Levy et al., 2021). Although further studies are needed, we suggest that the unique use of Cl^−^ as the counterion in opsins of ASO-II group, rather than a negatively charged amino acid, may be associated to their pH-sensitive light response and, ultimately, to their role in photosynthesis-related functions in symbiotic cnidarians.

It is widely accepted that opsins have evolved from a non-opsin GPCR (Feuda et al., 2012). Our finding of the native chloride counterion in opsins shows a possibility that in the first stage of the evolutionary process, the “primitive” opsins having Lys296 but no counterion residue could absorb not only UV light but also a wide range of wavelengths that extends into the visible region by embracing chloride ions. Namely, the chloride counterion system might have been a preliminary step in the evolution of amino acid counterions in animal opsins. Further empirical and bioinformatic studies are required for disentangling the evolutionary trajectories of the Schiff base – counterion system, including the chloride counterion.

## Materials and Methods

### Experimental design

We first identified and cloned opsins from a reef-building coral, *Acropora tenuis*, and then expressed opsins belong to the ASO-II group in mammalian cultured cells. We performed spectroscopic measurements of purified pigments of the opsins in different pH and Cl^−^ conditions to identify their effects on the acid dissociation constant of the protonated Schiff base of the opsins, leading to the determination of the counterion. Computational modelling and QM/MM calculations were also conducted to elucidate the retinylidene Schiff base environment in the dark state of Antho2a. Light-evoked Ca^2+^ responses were assessed by aequorin-based bioluminescent reporter assay to evaluate the G protein activation of Antho2a.

### Identification of *Acropora tenuis* opsins and phylogenetic tree inference

*Acropora tenuis* (Dana, 1846) is a common reef-building coral distributed throughout the Indo-Pacific Ocean. Candidate sequences of *A. tenuis* opsins were identified by homology search against public genome and transcriptome datasets (Shinzato et al., 2020; Voolstra et al., 2015) and their phylogenetic relationships to known opsins were inferred by subsequent phylogenetic tree reconstruction. We first conducted BLASTP and TBLASTN searches with an E-value cut-off of 10^−10^ using *Acropora palmata* Acropsin 1–3 (JQ966100 – JQ966102), two *Nematostella vectensis* opsins (BR000676 – BR000677), human rhodopsin (NM_000539), and squid rhodopsin (X70498) as queries. We aligned the collected opsin homologs and excluded sequences which did not contain a retinal-binding lysine residue (Lys296) in the seventh transmembrane helix. We modified the fragmented sequences by reference to the genome sequence of *A. tenuis* and opsin sequences of other *Acropora* species (*A. palmata* or *A. millepora*). The candidate sequences of *A. tenuis* opsins were combined with the representative opsin sequences. The final sequence set was aligned using MAFFT (Katoh and Standley, 2013) and trimmed by TrimAl (Capella-Gutiérrez et al., 2009) with the “*gappyout*” function. The ML tree was reconstructed using RAxML-NG v1.1.0 (Kozlov et al., 2019) assuming the LG + G4 model of protein evolution, which was selected by MoldelTest-NG v0.2.0 (Darriba et al., 2020). The ML branch supports were estimated with 1000 bootstrap replicates.

### Sample collection, total RNA extraction, and cDNA synthesis

Colonies of *A. tenuis* were collected from < 3 m depth on the fringing reef on Sesoko Island, Okinawa (N26°37.58′, E127°52.01′) and were maintained in flow-through aquaria at Sesoko Station (Tropical Biosphere Research Center, University of Ryukyus, Okinawa, Japan). Four days after spawning, motile larvae and small branches of adult colonies were preserved in RNAlater® Stabilization Solution (Thermo Fisher Scientific, MA, USA). Total RNAs were extracted from the larval and adult samples using TRIzol reagent (Thermo Fisher Scientific) or Sepasol-RNA I Super G (nacalai tesque, Kyoto, Japan) and purified using Qiagen RNeasy mini kits (Qiagen, Hilden, Germany) following the manufacturer’s protocol. cDNAs were synthesized from the total RNA by reverse transcription using High-Capacity cDNA Reverse Transcription kits (Thermo Fisher Scientific).

### Expression and purification of *Acropora tenuis* opsins

The coding regions of *A. tenuis* opsins were amplified by PCR with gene-specific primers and were tagged with the epitope sequence of the anti-bovine rhodopsin antibody rho 1D4 (ETSQVAPA) at their C-termini. Site-directed mutants were produced by overlap extension PCR using PrimeSTAR Max DNA Polymerase (TAKARA, Shiga, Japan) with site-specific primers and were also tagged with the 1D4 epitope sequence. The tagged cDNAs were inserted into the pUSRα vector (Kayada et al., 1995) digested with HindIII and EcoRI or the pMT vector (Ridge and Abdulaev, 2000) digested with EcoRI and NotI using In-Fusion HD cloning kit (TAKARA). The plasmids (15 µg per 100 mm culture dish) were transfected into COS-1 cells using the polyethyleneimine (PEI) transfection method as described previously (Obayashi et al., 2025; Sinha et al., 2014). The transfected cells were maintained for 24 h after transfection at 37°C under 5% CO_2_ and then 11-*cis* retinal was added to the medium (1 µL of 4 mM 11-*cis* retinal to 100 mm culture dish) following 25°C or 30°C incubation for another 24 h in the dark before collecting the cells. The reconstituted pigments were extracted from the cell membranes with 1% dodecyl β-D-maltoside (DDM, Dojindo, Kumamoto, Japan), 50 mM HEPES, and 140 mM NaCl (pH 6.5). The solubilized samples were mixed with 1D4-condjugated agarose beads overnight, and the mixture was transferred into Bio-Spin columns (Bio-Rad, Hercules, CA, USA) and washed in the buffer containing 0.02% DDM, 50 mM HEPES, and 140 mM NaCl (pH 6.5, buffer A). The purified pigments were eluted with buffer A containing 0.5 ∼ 1 mg/mL 1D4 peptide (custom peptide synthesis by GenScript Japan Inc., Tokyo, Japan). To obtain pigments in solutions of various anions (SO_42−_, Br^−^, and I^−^) other than Cl^−^, samples were prepared as described above and in the final step the mixture of solubilized samples and 1D4-agarose beads was washed with buffer A followed by the additional wash with buffers including different sodium salts of anions (0.02% DDM, each of 70 mM Na_2_SO_4_, 140 mM NaBr, or 140 mM NaI, and 50 mM HEPES). Then the pigments were eluted with the buffer including the appropriate sodium salt of anion containing 0.5 ∼ 1 mg/mL 1D4 peptide.

Alternatively, for some pigments that were unstable under the absence of Cl^−^, we quickly removed Cl^−^ by gel-filtration chromatography on PD MiniTrap desalting columns with Sephadex G-25 resin (Cytiva, Marlborough, MA, USA). The columns were first equilibrated with the buffer including 0.02% DDM, 70 mM Na_2_SO_4_, and 50 mM HEPES, 500 µL of samples were loaded onto the columns and eluted with the buffer. We collected 800 µL fractions and used them for subsequent spectroscopic analyses.

### UV-visible spectroscopy

Spectroscopic measurements were performed at 0°C using a V-750 UV-visible spectrophotometer (JASCO Corporation, Tokyo, Japan). The pH of the samples was adjusted with 100 mM CAPS including NaOH for alkaline conditions and 500 mM NaH_2_PO_4_ for acidic conditions. pH values were measured using a pH meter (B-211; HORIBA, Kyoto, Japan) immediately after each spectroscopic measurement. The concentration of Cl^−^ in the samples was adjusted by addition of different concentrations of NaCl solutions which were prepared in 70 mM Na_2_SO_4_ buffer (see above). A 100-W halogen lamp was equipped on the spectrophotometer and used to illuminate samples with a set of optical interference filters (420 nm or 500 nm, Toshiba) and cut-off filters (O-55 or O-56, AGC Techno Glass Co., Shizuoka, Japan). Absorption spectra of some UV-absorbing pigments were recorded using the V-750 UV-visible spectrophotometer, equipped with a 300-W xenon lamp (MAX-350; Asahi Spectra Co., Tokyo, Japan) that was used for illumination of samples in combination with a UV-transmitting filter (UTVAF-50S-36U, SIGMA KOKI, Tokyo, Japan).

### HPLC analysis

An HPLC analysis was carried out to analyze the conformations of retinal present in the purified pigments as described previously (Terakita et al., 1989), with some modifications. Briefly, 100 µL of purified pigments were mixed with 210 µL of cold 90% methanol which was stored in −20°C and 30 µL of 1 M hydroxylamine to convert retinal chromophore in a sample into retinal oxime. The retinal oxime was extracted with 700 µL of *n*-hexane. 200 µL of the extract were injected into a YMC-Pack SIL column (particle size 3 μm, 150 × 6.0 mm) and eluted with *n*-hexane containing 15% (v/v) ethyl acetate and 0.15% (v/v) ethanol at a flow rate of 1 mL/min.

### Bioluminescent reporter assays for Ca^2+^ measurements in cultured cells

Ca^2+^ levels in opsin-expressing cultured cells were assessed by an aequorin-based luminescent assay as described previously (Koyanagi et al., 2022). Briefly, the plasmid containing open reading frames of each opsin was transfected into HEK293S cells in 35 mm dishes by the PEI method with the aequorin plasmid obtained by introducing a reverse mutation A119D into the plasmid [pcDNA3.1+/mit-2mutAEQ] (Addgene no. 45539) (de la Fuente et al., 2012). The transfected HEK293S cells were incubated for ∼24 hours at 37°C under 5% CO_2_ with the addition of 0.2 µM/dish of 11-*cis* retinal 4 ∼ 5 hours after the transfection. Before the luminescence measurements, the culture medium was replaced with a medium containing Coelenterazine *h* (pH 7.0), and the cells were incubated to equilibrate with the media at 25°C for at least 2 hours. Dishes of cells were then stimulated with light and luminescence values were recorded using GloMax 20/20n Luminometer (Promega). A green (495 nm) LED light (color: “Cyan”, Ex-DHC; BioTools Inc., Gunma, Japan) and arrayed LEDs on a board with spectral emission peaks at 390 nm and 510 nm (SPL-25-CC; REVOX Inc., Kanagawa, Japan) were used as light sources.

### Computational modelling and QM/MM calculations

The three-dimensional structure of Antho2a was predicted from the primary amino acid sequence using AlphaFold2 (Jumper et al., 2021). The 11-*cis* retinal chromophore linked to the protonated Schiff base was incorporated into the AlphaFold model using as a template the high-resolution structure of bovine rhodopsin solved by time-resolved serial femtosecond X-ray crystallography (Gruhl et al., 2023). A Cl^−^ anion was initially placed in close proximity to the retinal protonated Schiff base, as observed in the microbial chloride-pump halorhodopsin (Mous et al., 2022). Water molecules were added using HomolWat (Mayol et al., 2020). We determined the p*K_a_* values of the titratable amino acid residues at pH 6.5 using the PROPKA program (Olsson et al., 2011; Søndergaard et al., 2011) and subsequently, the protein was protonated using the tleap program in the AMBER software package (Case et al., 2016). The geometry of this initial model was first relaxed by molecular mechanics energy minimization with the Amber ff14SB force field (Maier et al., 2015) using steepest descent for 10,000 steps before switching to a conjugate gradient minimiser for an additional 10,000 steps. During energy minimization, a positional restraint of 10 kcal mol^-1^Å^-2^ was applied to all atoms, including hydrogens. The SHAKE algorithm (Ryckaert et al., 1977) was used to constrain the motion of bonds involving hydrogen. Finally, the geometry of the system was optimized using hybrid quantum mechanics/molecular mechanics (QM/MM) calculations without considering any external environment and with the backbone of the protein frozen (Mroginski et al., 2021; Senn and Thiel, 2009). The QM part consists of the retinal chromophore linked to the lysine sidechain cut between the Cδ and Cε atoms forming the protonated Schiff base, along with Cl^−^, Glu292 and Ser186. The retinal-binding pocket also contains predicted water molecules (modelled based on homologous GPCR structures) close to the Schiff base and the chloride ion which were not included in the QM region. The hydrogen link atom (HLA) scheme was used at the QM/MM boundary. The QM part was treated using the BP86-D3 (BJ) functional (Becke, 1988; Grimme et al., 2011) in conjunction with the cc-pVDZ basis set (Dunning, 1989) and the def2/J auxiliary basis set for the resolution of identity (RI) (Weigend, 2008). The Chain of Spheres exchange (COSX) algorithm was utilized in combination with the resolution of identity for the Coulomb term (RI-J). The rest of the protein was treated with the Amber ff14SB force field. Water molecules were treated with the TIP3P model (Jorgensen et al., 1983). The QM/MM calculations were performed using the quantum chemistry program Orca 5.0.2 (Neese, 2022) interfaced with the DL_POLY module of the ChemShell 3.7.1 software package (Metz et al., 2014; Sherwood et al., 2003). The optimized ground state geometries and partial charges were used to calculate the vertical excitation energies using the simplified time-dependent density functional theory (sTD-DFT) (Bannwarth and Grimme, 2014; Runge and Gross, 1984) at the CAM-B3LYP/cc-pVTZ level of theory (Dunning, 1989; Yanai et al., 2004) using the Orca program. The excitation energies were also calculated using the RI-ADC(2) method (Hättig, 2005) with frozen core orbitals and the cc-pVTZ basis set in association with the corresponding auxiliary basis by utilizing the Turbomole 7.5.1 program package (Furche et al., 2014). The three-dimensional models were visualized using the molecular graphics program PyMOL 2.5.5.

## Acknowledgements

We thank Dr. Robert S. Molday (University of British Columbia) for kindly supplying rho 1D4-producing hybridoma. We thank Dr. Masayuki Hatta (Ochanomizu University) for coral sampling. We are also grateful to Dr. Hisao Tsukamoto for technical guidance on the transfection of COS-1 cells. This work was supported by the Japanese Ministry of Education, Culture, Sports, Science and Technology Grants-in-Aid for Scientific Research 23H02516 (to AT), 22H02663 (to MK) and JP20J01841 (YS); and Japan Science and Technology Agency (JST) Core Research for Evolutional Science and Technology (CREST) Grant JPMJCR1753 (to AT). YS was supported by Grant-in-Aid for JSPS Fellows. This work has also been supported by the Swiss National Science Foundation (project grant 192780 to XD) and by the European Union’s Horizon 2020 research and innovation programme (grant agreement 951644 to GFXS and Marie Skłodowska-Curie grant agreement 884104 (PSI-FELLOW-III-3i) to SS). We finally thank the members of the High-Performance Computing and Emerging Technologies (HPCE) group at the Paul Scherrer Institute for technical support and assistance with high-performance computing.

## Author contributions

YS, MK, and AT conceptualized the study; YS, MK, GFXS, XD, and AT designed the study; YS, TS, YK, and MI performed the experiments and analyses; SS and XD performed the computational modelling; YS and AT wrote the original draft with inputs from SS and XD; All authors reviewed the manuscript and gave final approval for the manuscript.

## Competing interests

The authors declare that they have no competing interests.

## Data and materials availability

The cDNA sequences of Acropora tenuis opsins in this paper are available in GenBank: eight Cnidopsins (accession no. LC844924–LC844931), seven opsins in the ASO-II group (accession no. LC844932–LC844938), and one opsin in the ASO-I group (accession no. LC844939). The structural models of wild type Antho2a with a neutral or charged Glu292 and the Antho2a E292A mutant are available in Zenodo (10.5281/zenodo.15064942). All data needed to evaluate the conclusion in this paper are present in the paper and the Supplementary Materials. Additional data related to this paper may be requested from the authors.

---

**Fig. S1.**
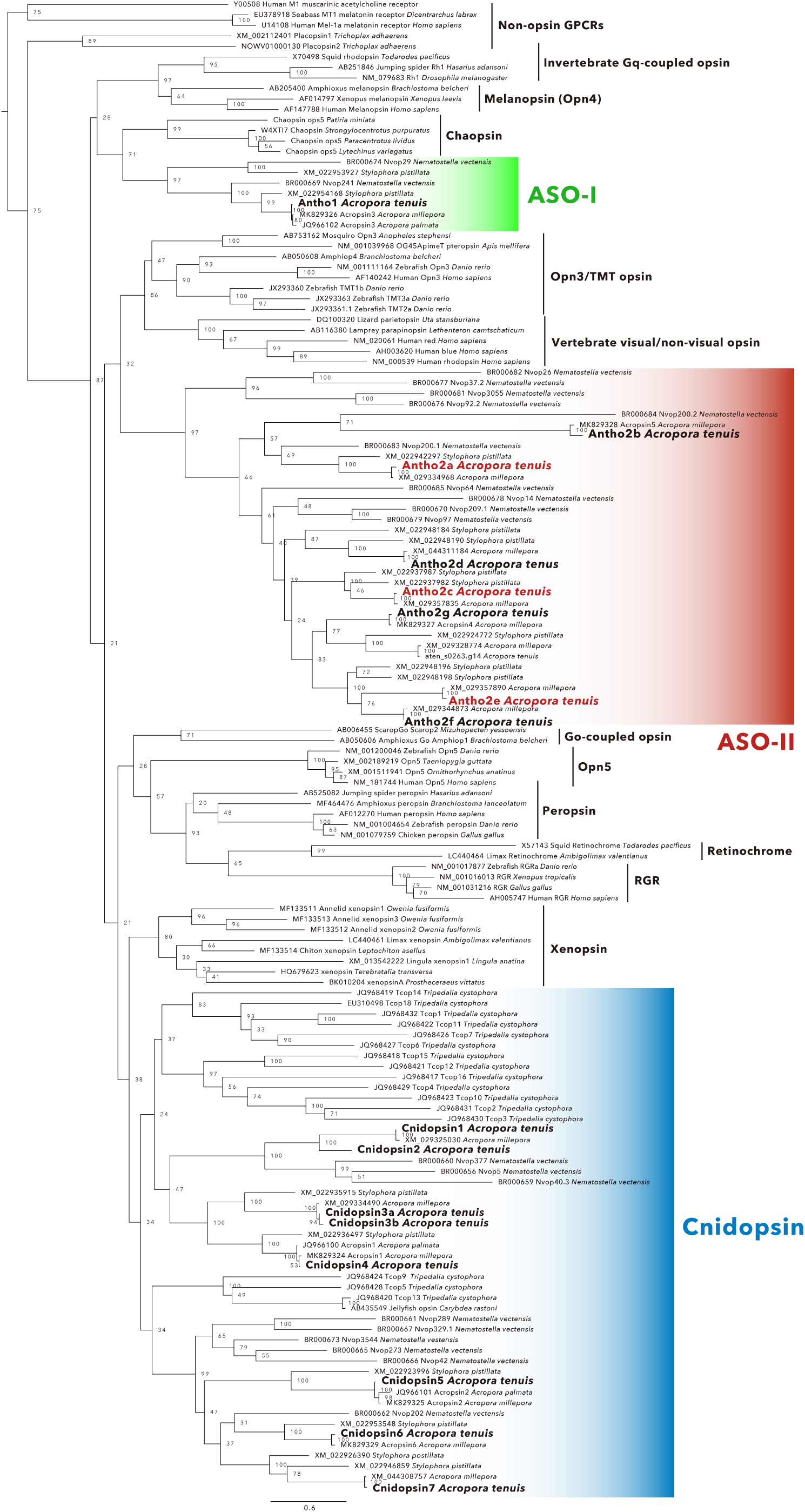
Maximum-likelihood (ML) tree of animal opsins, with non-opsin GPCRs included as an outgroup. (a simplified version of the ML tree is shown in Fig. 1A). The tree includes opsins belonging to the eight main groups (see main text) as well as of opsins from the recently identified subgroups xenopsins and chaopsins (Ramirez MD et al. *Genome Biol Evol* **8**:3640–3652, 2016). The sixteen *A. tenuis* opsins (eight opsins in the cnidopsin group, one opsin in the ASO-I group, and seven opsins in the ASO-II group) that were identified in this study are shown in bold. The three opsins in the ASO-II group for which absorption spectra were successfully obtained are highlighted in red. Numbers at nodes represent support values for the ML branch estimated by 1000 bootstrap samplings (≥ 70% are indicated). Scale bar = 0.6 substitutions per site.

**Fig. S2.**
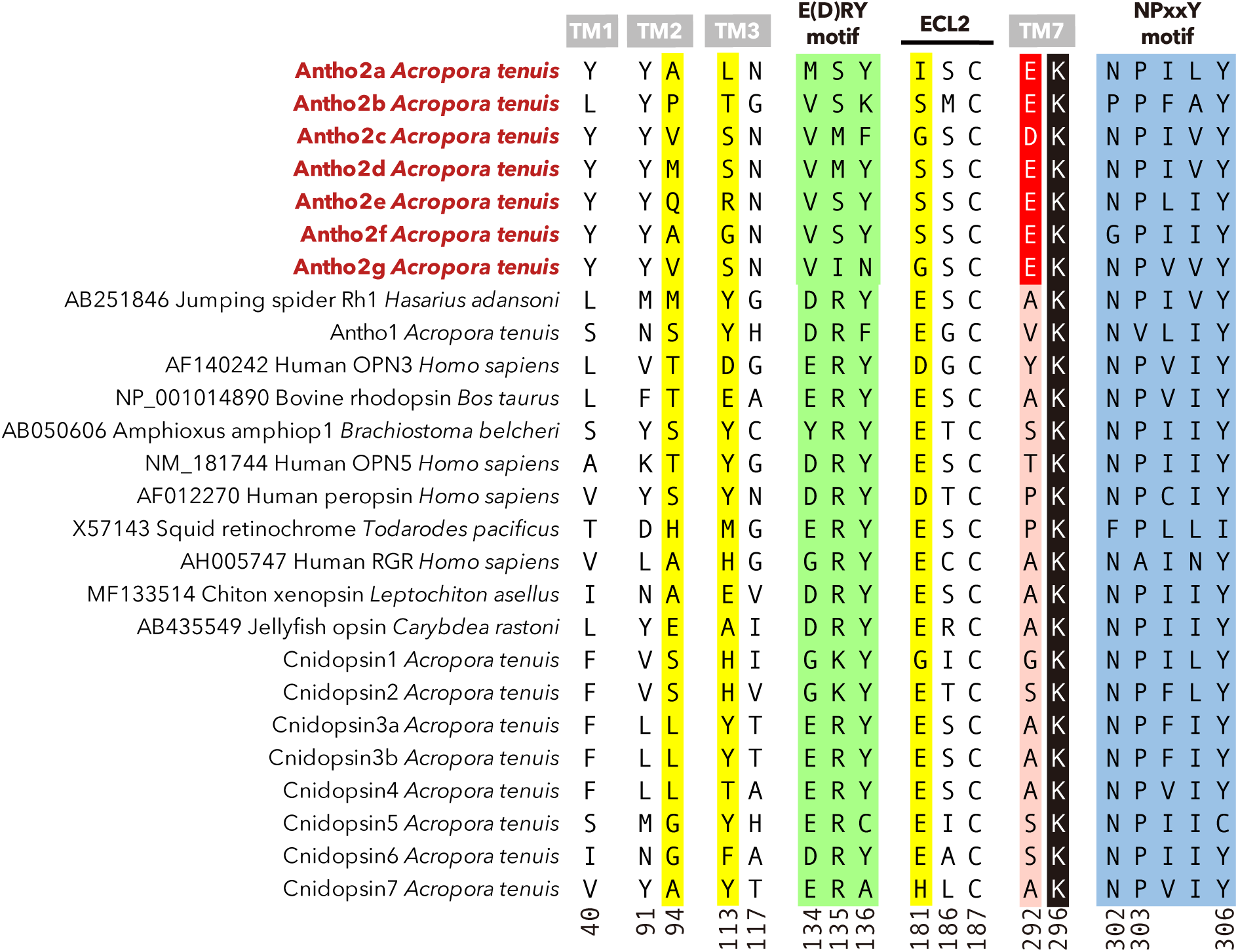
Key residues in opsins of the ASO-II group and other animal opsins. Residues near the Schiff base were selected with reference to the crystal structure of bovine rhodopsin (PDB ID: 1U19) and the homology model of Antho2a (shown in Fig. 4). Position 292 contains an acidic residue (D/E) only in opsins of the ASO-II group (red). In addition, functionally important residues (such as the retinal binding Lys296 (black), the three established counterion sites (yellow), and two highly conserved motifs in Class A GPCRs, E(D)RY on TM3 (green) and NPxxY on TM7 (blue)) are also shown. Residues are numbered according to bovine rhodopsin.

**Fig. S3.**
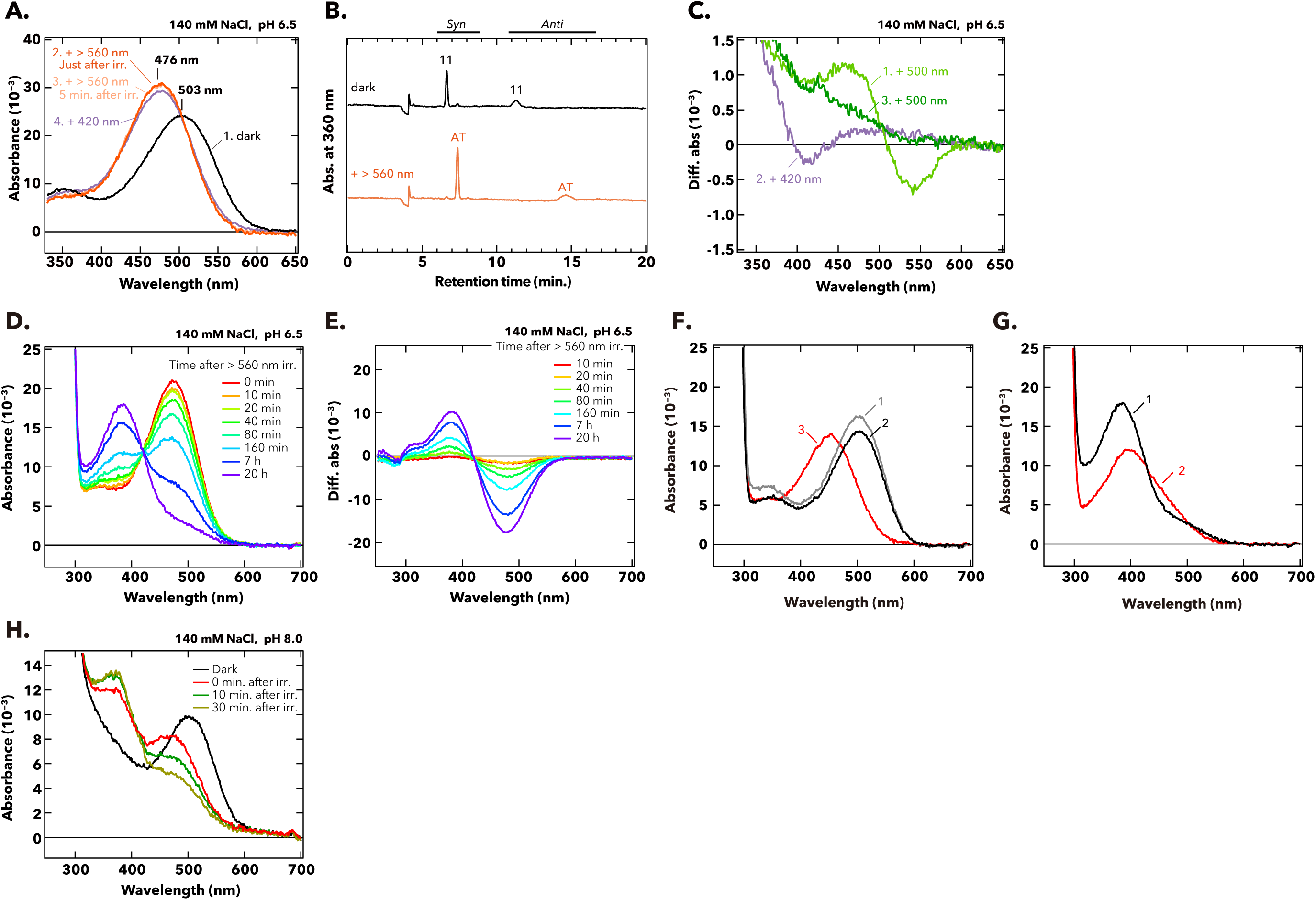
Photo- and thermal reactions of wild type Antho2a. (A) Absorption spectra of purified wild type Antho2a were measured at 0°C in the dark (curve 1, black), after orange light irradiation (> 560 nm, curve 2 and curve 3, deep and pale orange), and after subsequent violet light irradiation (420 nm, curve 4, purple). Irradiation of wild type Antho2a with orange light shifted the absorption maximum from 503 nm in the dark (curve 1, black) to 476 nm in the photoproduct (curve 2 and curve 3, orange). Upon subsequent irradiation with 420 nm light (while the photoproduct was stable), the λ_max_ of the photoproduct stayed at 476 nm, with only a slight decrease of the peak absorbance (curve 4, purple) possibly resulting from degradation of the photoproducts upon light irradiation. (B) The configuration of retinal in wild type Antho2a before (black) and after (orange) irradiation with orange light (> 560 nm) was analyzed with HPLC. Retinal was extracted in its oxime form. AT, all-*trans* retinal; 11, 11-*cis* retinal. (C) Difference spectra calculated from the absorption spectra recorded before and after sequential irradiations with 500 nm and 420 nm light for crude extracts of detergent-solubilized cell membranes expressing wild type Antho2a. Curve 1: Difference spectrum of after minus before 500 nm irradiation, showing a blue-shifted spectral change indicative of photoproduct formation. Curves 2 and 3: Difference spectra of after minus before subsequent 420-nm light irradiation (curve 2), and after minus before a second 500 nm light irradiation, following the 420 nm irradiation (curve 3). Neither condition resulted in spectral changes consistent with regeneration of the dark state, indicating that Antho2a is not bistable under these conditions. (D) Changes in the absorption spectra of the purified pigment of wild type Antho2a after irradiation with orange (>560 nm) light (with the sample kept in the dark at 0°C). Each colored curve corresponds to a different incubation time after the light irradiation. (E) Difference spectra obtained by subtracting the spectrum of wild type Antho2a in the dark from the spectra measured at different time points after irradiation (shown in D). (F, G) Acid denaturation of pigments before (dark state, F) and after light irradiation (G). (F) Absorption spectra of wild type Antho2a in the dark were measured immediately after sample preparation at pH 6.5 (curve 1), and after incubation over night at 0°C (curve 2, pH 6.5) with subsequent addition of HCl to pH 1.9 (curve 3). (G) Absorption spectra of wild type Antho2a incubated for 20 hours at 0°C following irradiation with orange light (> 560 nm), measured at pH 6.5 (curve 1) and after acidification to pH 1.9 (curve 2). When retinal binds to opsin via a Schiff base (protonated or deprotonated), acid denaturation traps the chromophore as a protonated Schiff base, exhibiting absorption at λ_max_ ∼440 nm. Acid denaturation of irradiated Antho2a (incubated for 20 hours post-irradiation) did not yield a product with λ_max_ at ∼440 nm (G, curve 2), whereas the dark state pigment after acidification displayed absorption at ∼440 nm (F, curve 3). These results indicate that Antho2a gradually releases retinal after light irradiation. (H) Absorption spectra of purified wild type Antho2a at pH 8.0, measured at 0°C before (dark state; curve 1, black) and 0, 10, and 30 minutes after irradiation with orange light (> 560 nm; red, green, and yellow curves, respectively).

**Fig. S4.**
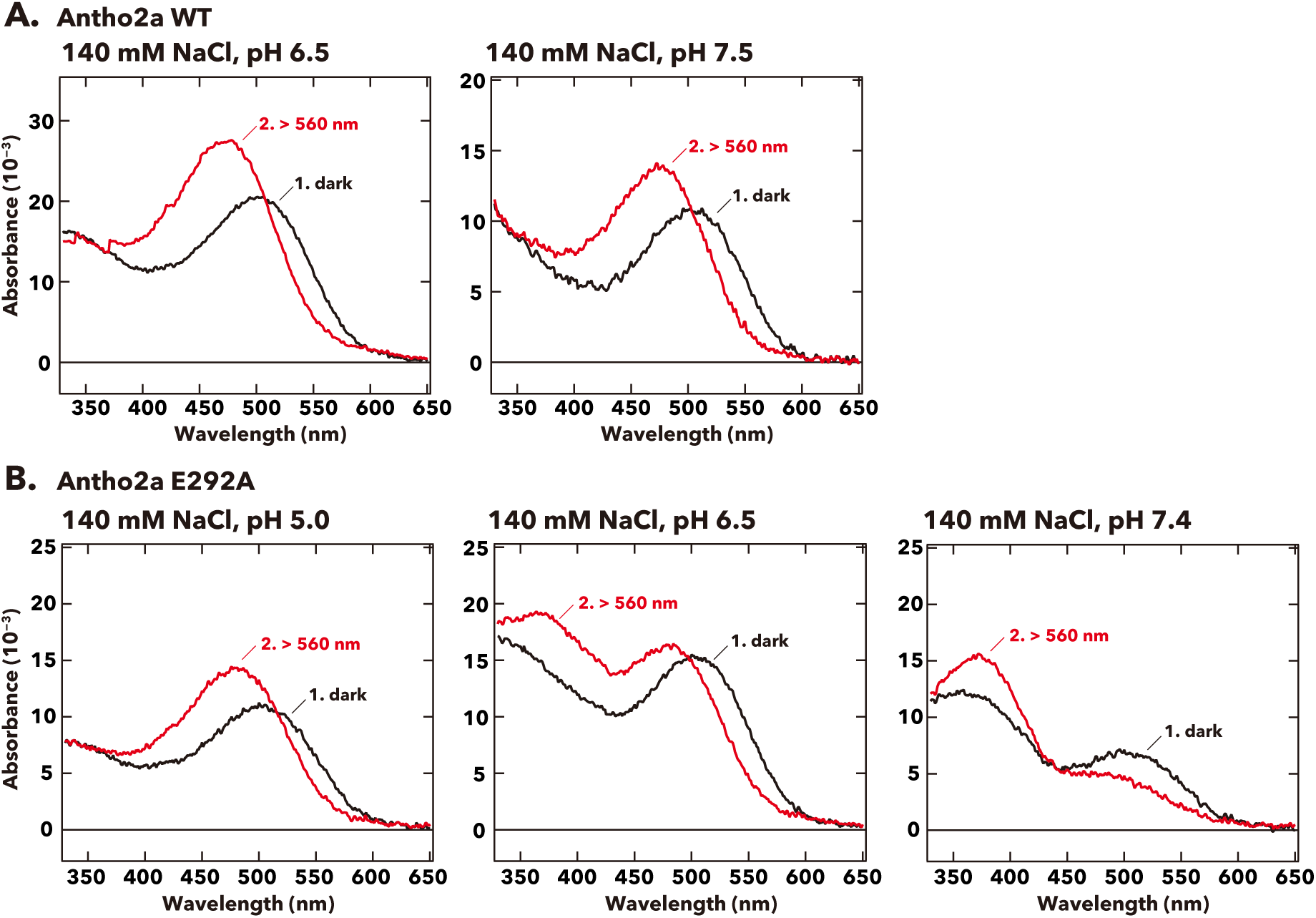
(A, B) Effect of pH on the absorption spectra of the dark state and photoproduct of (A) wild type Antho2a and (B) the E292A mutant at 140 mM NaCl. Each graph shows spectra before (curves 1, black) and after (curves 2, red) irradiation with orange (> 560 nm) light.

**Fig. S5.**
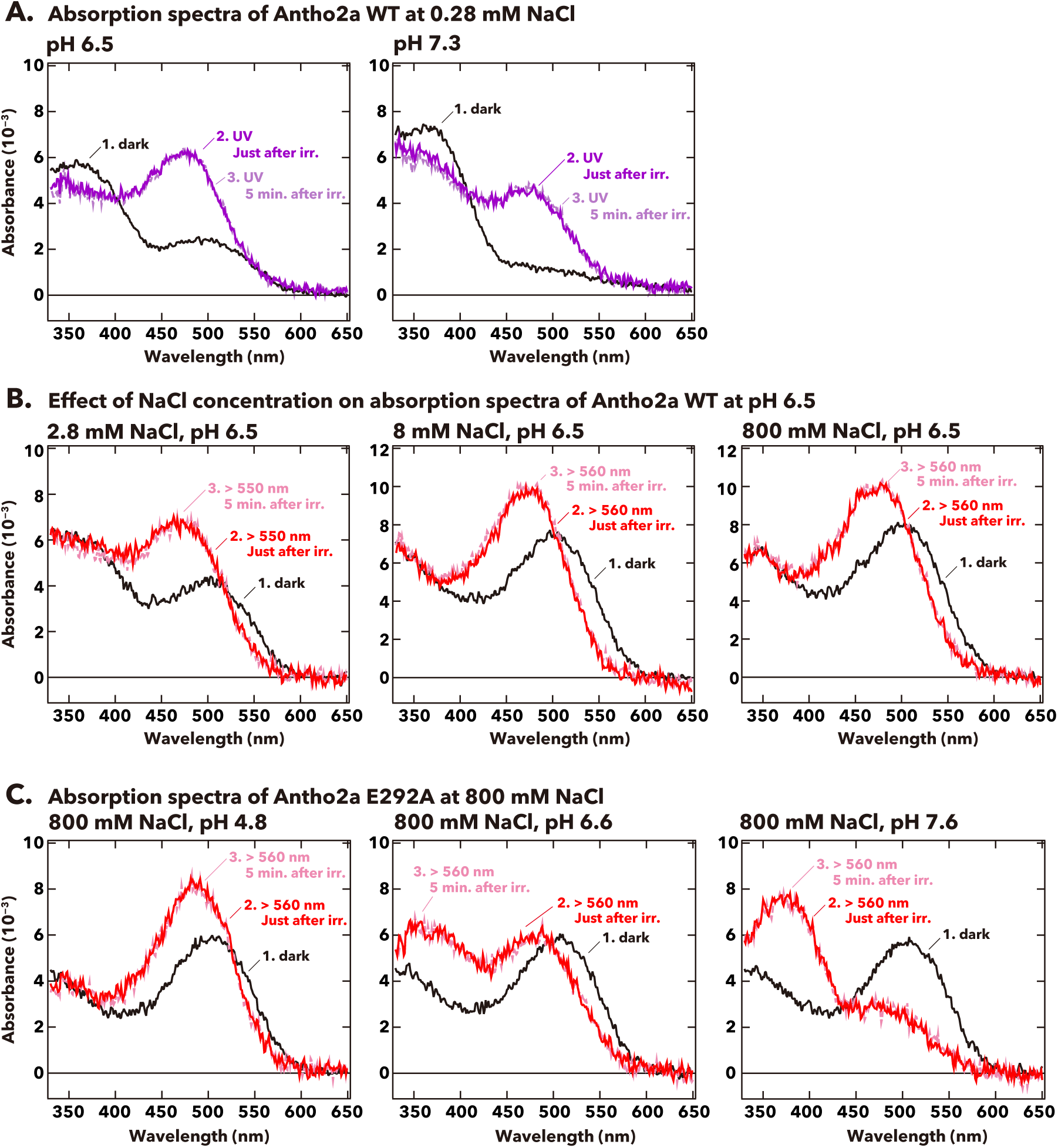
Effect of pH and NaCl concentration on absorption spectra of the dark states and photoproducts of Antho2a. (A) Absorption spectra of wild type Antho2a at 0.28 mM NaCl under different pH conditions (pH 6.5 and pH 7.3). (B) Absorption spectra of wild type Antho2a at pH 6.5 with varying NaCl concentrations (2.8 mM, 8 mM, and 800 mM). (C) Absorption spectra of the E292A mutant of Antho2a at 800 mM NaCl and different pH conditions (pH 4.8, pH 6.6, and pH 7.6). Each graph shows spectra before (curves 1, black) and after (curves 2, deep purple/red, and 3, pale purple/pink) irradiation with UV light (< 410 nm, A) or orange light (> 550 nm or > 560 nm, B and C). Curves 2 and 3 in each graph represent, respectively, the first measurement (immediately after irradiation) and a subsequent measurement (< 5 min. after irradiation). The minimal change in the absorption spectra over this time scale indicates that the spectra remain stable after irradiation.

**Fig. S6.**
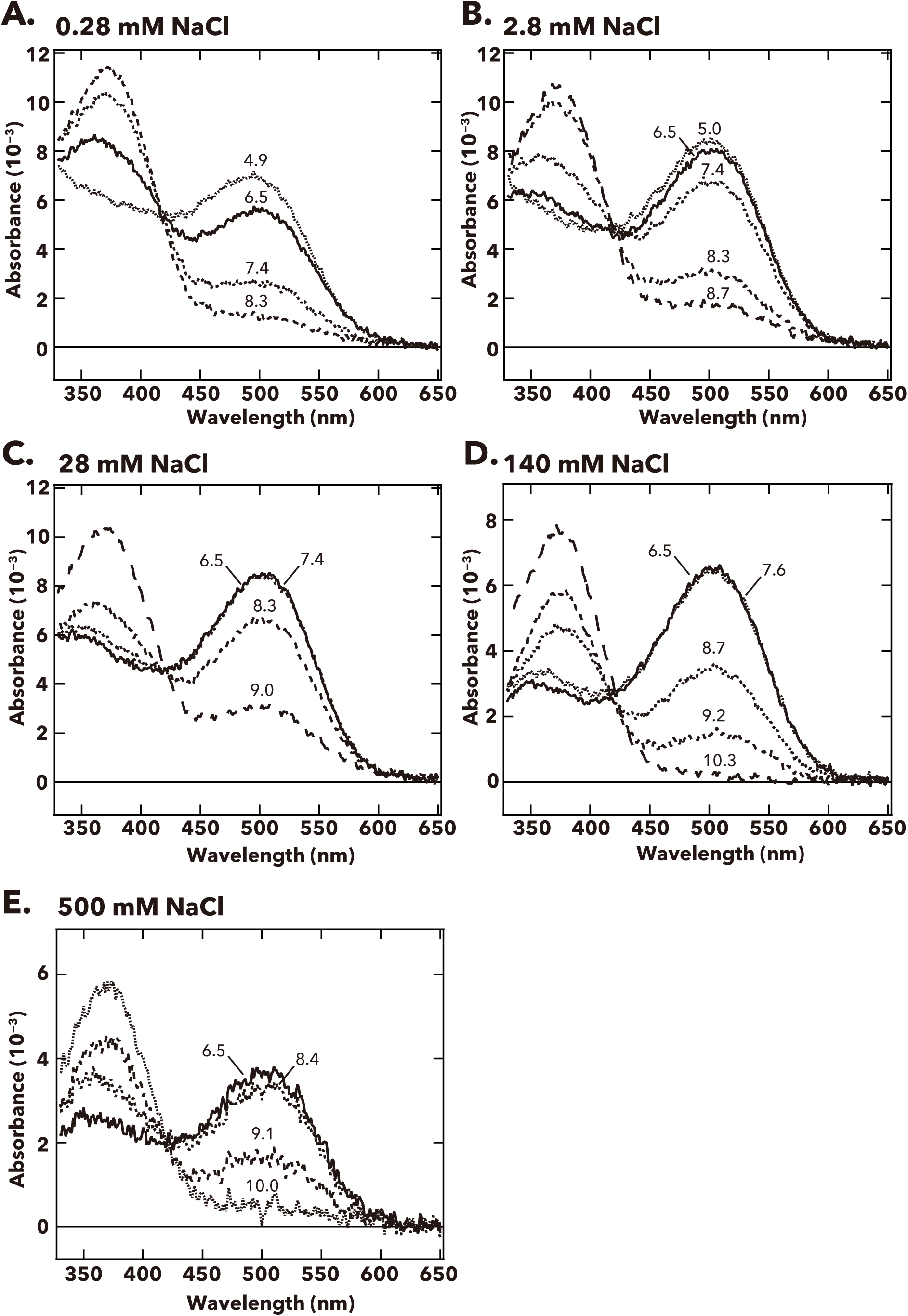
pH-dependent changes in the absorption spectra of Antho2a WT at (A) 0.28 mM, (B) 2.8 mM, (C) 28 mM, (D) 140 mM, and (E) 500 mM Cl^−^ at 0°C. The pH values of the solution, measured right after each spectroscopic measurement, are indicated on the corresponding curves in each graph.

**Fig. S7.**
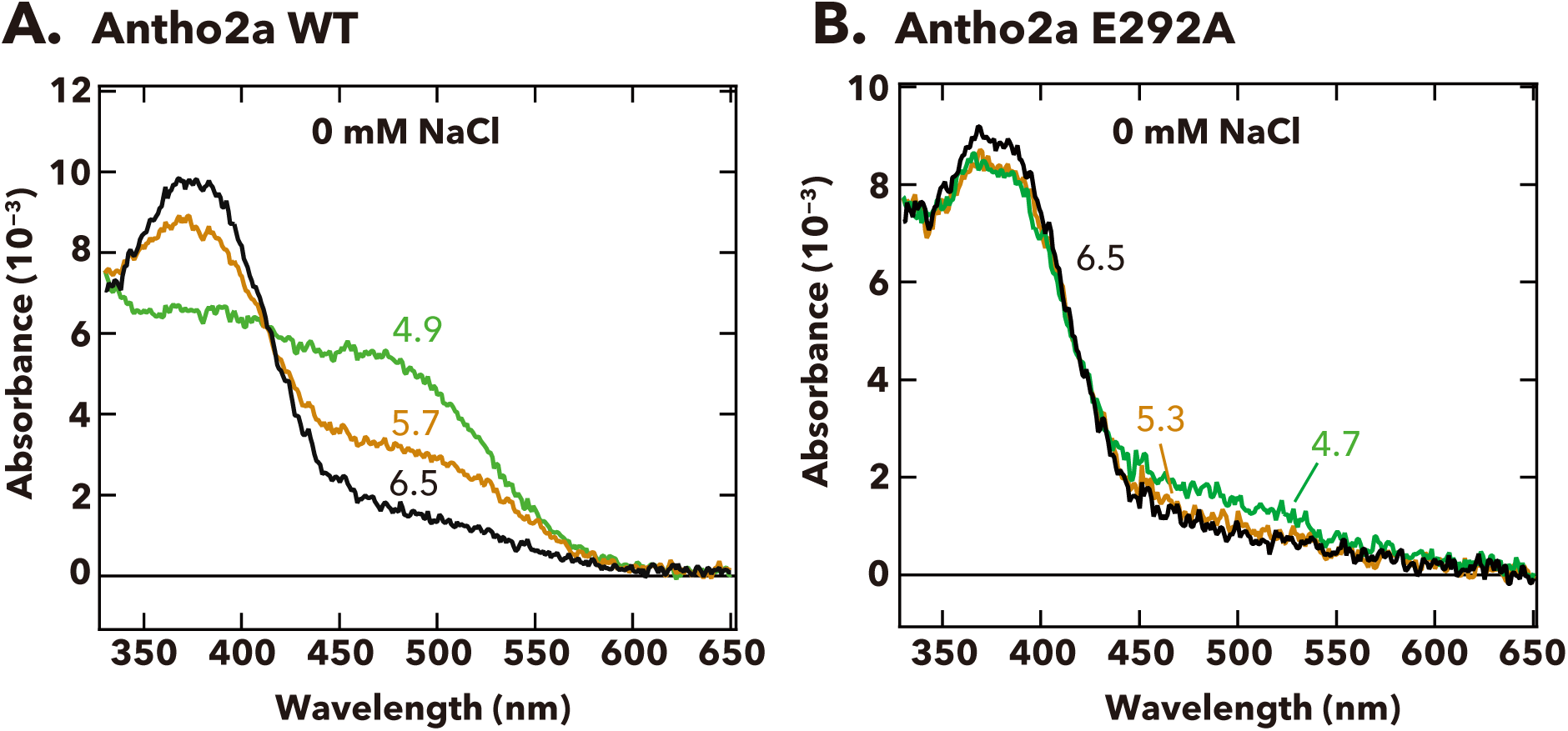
pH-dependent changes in the absorption spectra of Antho2a WT and Antho2a E292A at 0 mM NaCl (containing 70 mM Na_2_SO_4_) at 0°C. The pH values at which the absorption spectra were measured are indicated on the corresponding curves.

**Fig. S8.**
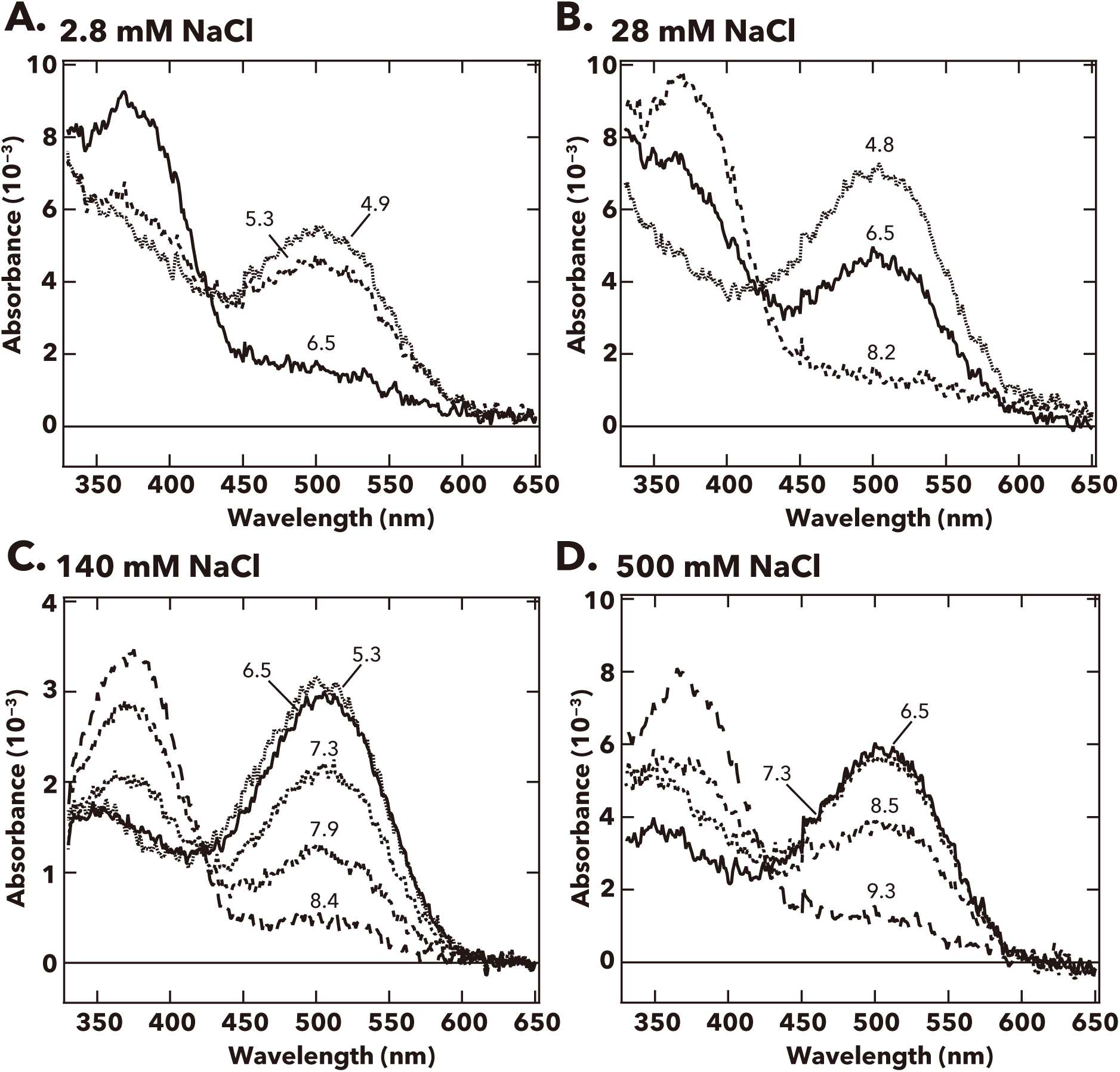
pH-dependent changes in the absorption spectra of Antho2a E292A at (A) 2.8 mM, (B) 28 mM, (C) 140 mM, and (D) 500 mM Cl^−^ at 0°C. The pH values are indicated on the corresponding curves in each graph.

**Fig. S9.**
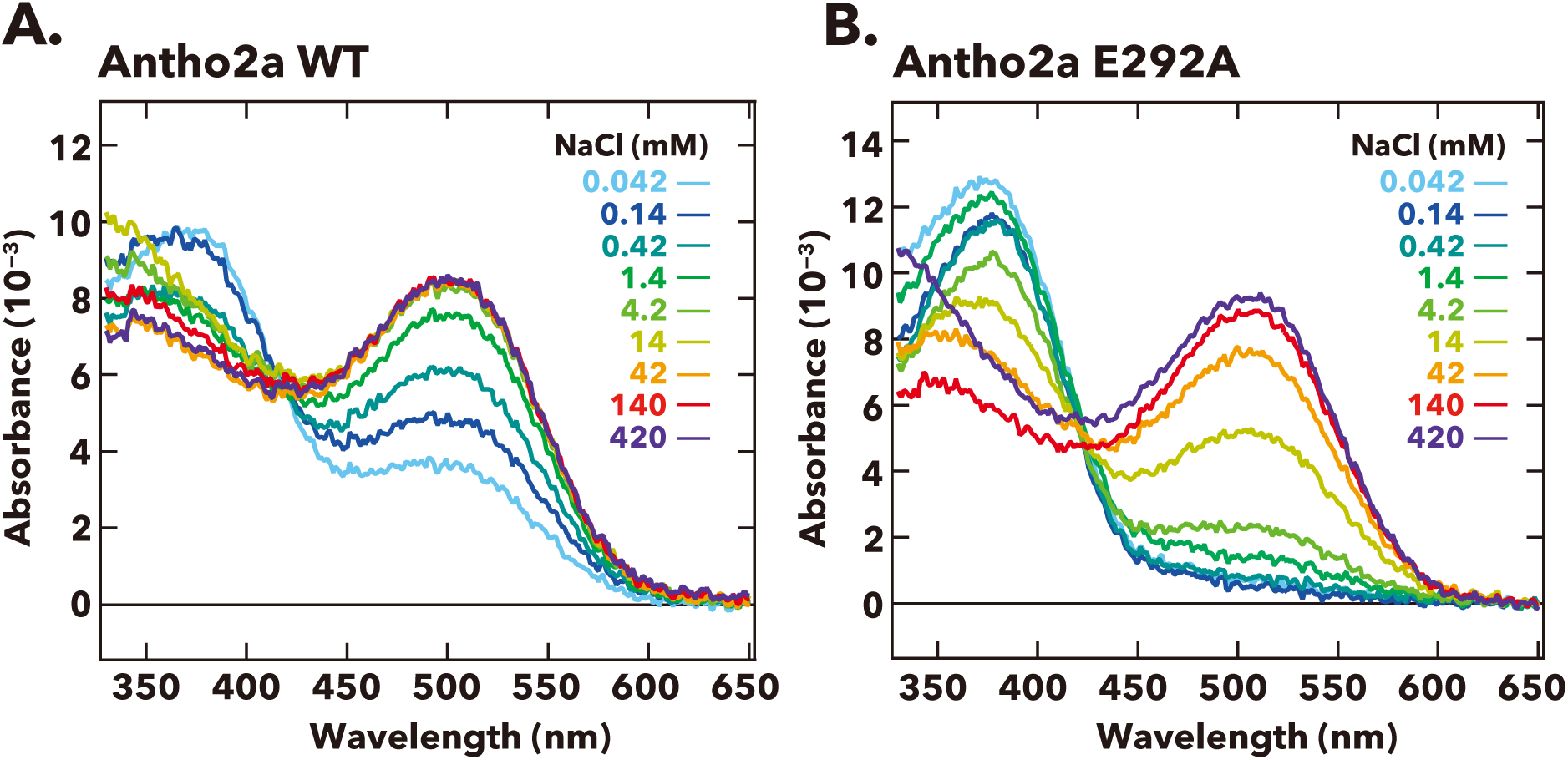
Absorption spectra of (A) wild type Antho2a and (B) the Antho2a E292A mutant under different Cl^−^ concentrations at pH 6.5 and 0°C. Each color indicates a different concentration of Cl^−^.

**Fig. S10.**
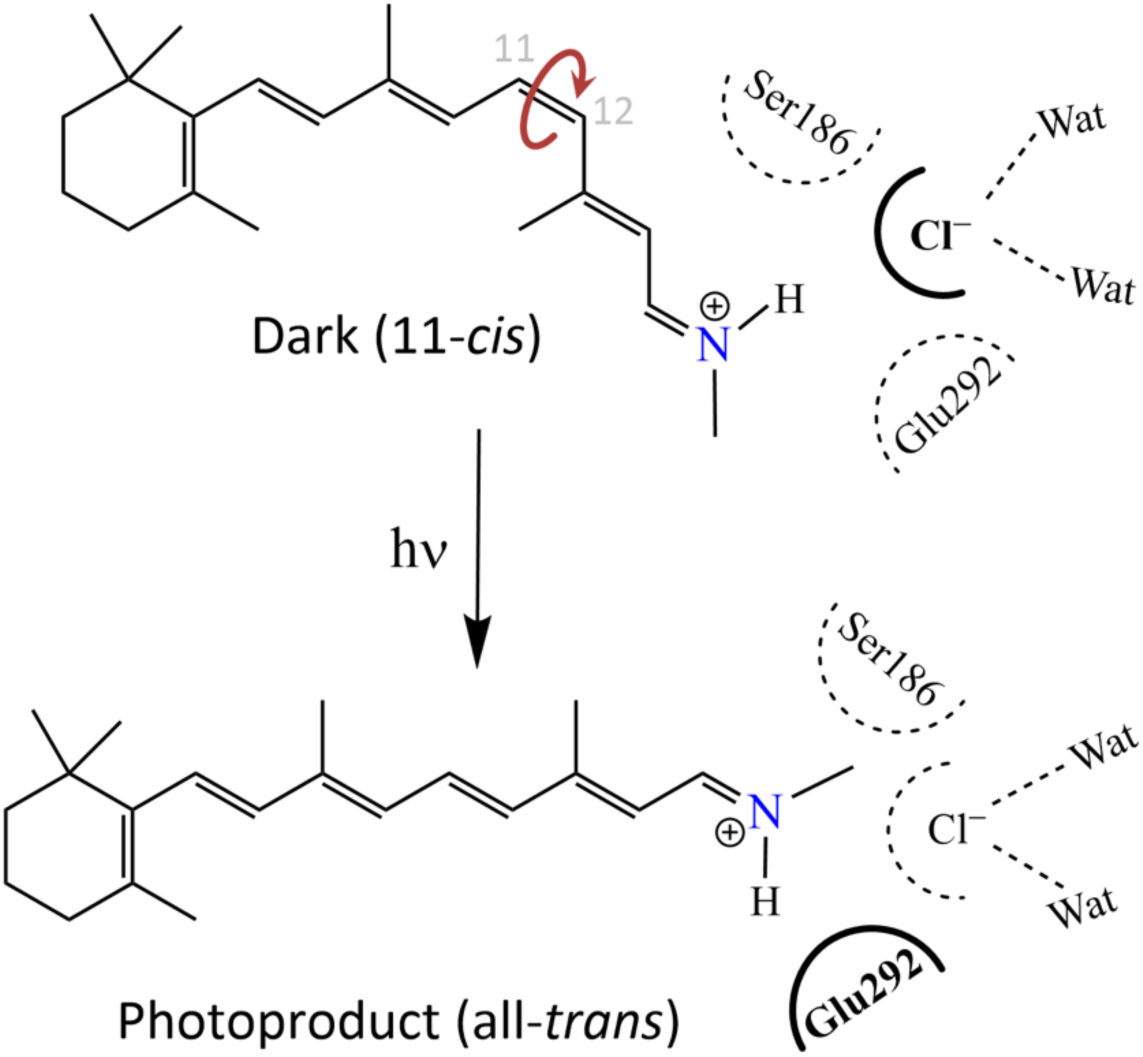
Schematic drawing of the environment of the protonated Schiff base depicting the counterion switch from a chloride ion (top) to Glu292 (bottom) upon light activation.

**Fig. S11.**
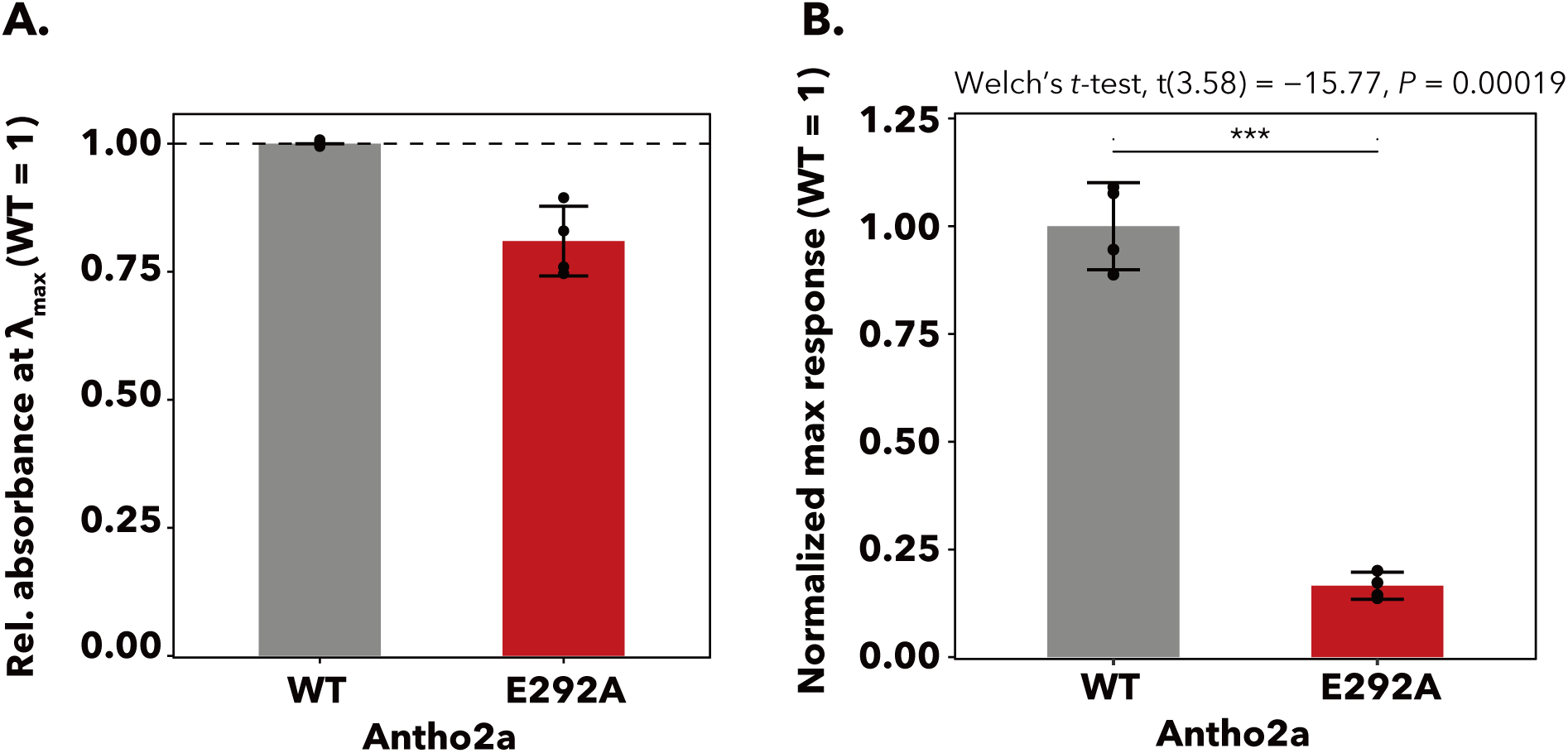
Expression levels and normalized Ca^2+^ responses of wild type and E292A Antho2a. (A) The relative absorbances at λ_max_ (± 5 nm) of wild type (grey) and E292A (red) Antho2a are used as indicators of expression levels. The proteins were expressed and purified under identical conditions, and absorbance values of the E292A mutant were normalized to those of the wild type. Spectra were measured at 0°C, pH 6.5, and 140 mM NaCl. Each bar represents mean ± S.E.M. of n = 4 replicates from separate transfections in cultured cells. (B) Normalized maximum Ca^2+^ responses of HEK293S cells expressing wild-type Antho2a and the E292A mutant after irradiation with green light (510 nm). Responses were normalized to the expression levels shown in panel A. Each bar represents mean ± S.E.M. of n = 4 replicates from separate transfections in cultured cells. Statistical evaluation of the normalized Ca^2+^ responses for wild type Antho2a versus the E292A mutant was conducted using a Welch’s *t*-test (two-sided). \*\*\**P* < 0.001.

**Table S1.** Vertical excitation energies (Δ*E_calc_*) and oscillator strengths (*f*) computed by quantum mechanics/molecular mechanics (QM/MM) calculations using different QM methods with the cc-pVTZ basis set.

| QM Region | sTD-DFT CAM-B3LYP |  | ADC(2) |  |
| --- | --- | --- | --- | --- |
| | $\Delta E_{calc}$<br>nm (eV) | $f$ | $\Delta E_{calc}$<br>nm (eV) | $f$ |
| RET+Lyr296+Cl+Ser186+**Glu292 (deprotonated)** | 415 (2.99) | 1.54 | 374 (3.32) | 1.64 |
| RET+Lyr296+Cl+Ser186+**Glu292 (protonated)** | 503 (2.47) | 1.19 | 416 (2.98) | 1.43 |
| RET+Lyr296+Cl+Ser186+**Ala292** | 499 (2.49) | 1.19 | 426 (2.91) | 1.36 |

## Notes

### Competing Interest Statement

The authors have declared no competing interest.

### Summary of Updates

We revised method section (L435-L436) to cite the prior work mentioned by the reviewer and also revised acknowledgement (L537).

